# Reconstructing Sordida subcomplex (Hemiptera, Reduviidae, Triatominae) phylogeny across species distribution range

**DOI:** 10.1101/2025.01.30.635797

**Authors:** Gabriela Burgueño-Rodríguez, Julieta Nattero, Néstor Ríos, Romina Valeria Piccinali, Ana L. Carbajal-de-la-Fuente, Francisco Panzera, Catarina Macedo Lopes, Patricia A. Lobbia, Antonieta Rojas de Arias, María J. Cavallo, Claudia S. Rodríguez, Pedro Lorite, María C. Vega-Gómez, Miriam Rolon, Sebastián Pita

## Abstract

The conformation of the Sordida subcomplex has been a topic of prolonged debate, with diverse methodological approaches employed to discern its constituent species. Up to now, *Triatoma sordida*, *T. garciabesi* and *T. rosai* comprise part of this subcomplex. Distinguishing and identifying these three species pose significant challenges due to their pronounced morphological similarity, overlapping distributions, and presence of natural hybrids. This study aims to uncover the genetic diversity and geographic spread of these three species by analyzing a mitochondrial *cytochrome b* gene fragment and complementing it with chromosomal studies across natural populations from an extensive geographical range, including Argentina, Bolivia, Brazil, and Paraguay. Phylogenetic analyses revealed genetic distances that suggest the presence of at least six putative species, rather than the three currently recognized. The present findings underscore the potency and significance of molecular analyses from natural populations for species identification and highlight the limitations of morphology in classifying Triatominae species.

## INTRODUCTION

The Triatominae subfamily (Hemiptera, Reduviidae) comprises over 154 species of blood-sucking insects that vary in various aspects of their biology, particularly in their significance as vectors of Chagas disease (Lent and Wygodzinsky 1979, Téllez-Rendón *et al*. 2023, Zhao *et al*. 2023, Campos *et al*. 2024). Among the 18 genera comprising the subfamily, the genus *Triatoma* Laporte, 1832 stands out as the largest with more than 80, categorized into eight complexes and 15 subcomplexes (Monteiro *et al*. 2018, Campos *et al*. 2024). One of these is the Sordida subcomplex, where so far six species have been formally recognized: *Triatoma sordida* (Stål, 1859), *Triatoma garciabesi* (Carcavallo, Cichero, Martínez, Prosen & Ronderos, 1967), *Triatoma rosai* (Alevi, Oliveira, Garcia, Cristal, Delgado, Bittinelli, Reis, Ravazi, Oliveira, Galvão, Azeredo-Oliveira, Madeira, 2020), *Triatoma jurbergi* (Carcavallo, Galvão and Lent, 1998), *Triatoma matogrossensis* (Leite & Barbosa 1953) and *Triatoma vandae* (Carcavallo, Jurberg, Rocha, Galvão, Noireau, Lent, 2002). These six species (*T. rosai* was referred as *T. sordida* Argentina) were grouped within the Sordida subcomplex through chromosomal markers (Pita *et al*. 2016), and later confirmed through analysis of nuclear and mitochondrial DNA sequences (Monteiro *et al*. 2018, Belintani *et al*. 2020). The initial inclusion of *Triatoma guasayana* (Wygodzinsky and Abalos, 1949) and *Triatoma patagonica* (Del Ponte, 1929) in this subcomplex, based on morphological similarities (Carcavallo *et al*. 2000, Schofield and Galvão 2009), has been discarded by numerous chromosomal, molecular and chemical evidences (Pita *et al*. 2016, Monteiro *et al*. 2018, Belintani *et al*. 2020, Moriconi *et al*. 2022). Nevertheless, several authors based on mitochondrial and nuclear sequences, grouping T. guasayana within the Sordida subcomplex (Alevi *et al*. 2020; Kieran *et al*. 2021).

Until the revision made by Lent and Wygodzinsky (1979), *T. sordida* constituted a widely distributed species along Argentina, Bolivia, Brazil, Paraguay and Uruguay, showing variability in external coloration and body size. This species, frequently infected with the parasite *Trypanosoma cruzi* (Chagas, 1909), showed populations colonizing the domicile and peridomicile in some areas, while in other regions, it was observed in a great diversity of wild habitats (Lent and Wygodzinsky 1979). Since the 1950s, variations in external coloration and body size have been described among *T. sordida* from northeastern Argentina (large and domestic individuals with light color) and individuals from northwestern Argentina (small and sylvatic dark specimens) (Abalos and Wygodzinsky 1951, Usinger *et al*. 1966). Crossbreeding experiments between the two Argentinian groups demonstrated complete fertility for a minimum of two generations, although the data were not published by Wygodzinsky and Abalos (Usinger *et al*. 1966). Actis *et al*. (1964) described variations in the electrophoretic profiles of the hemolymph proteins between sylvatic individuals from northwest Argentina (smaller and darker) and domestic individuals from Brazil. In 1967, based on chromatic and morphological differences with domestic *T. sordida*, the northwestern Argentinian individuals were described as a new species named *T. garciabesi* (Carcavallo *et al*. 1967). But this species was subsequently synonymized with *T. sordida* by Lent and Wygodzinsky (1979). Later, Panzera *et al*. (1997) identified three isoenzymatic groups within *T. sordida*: the Northwest and Northeast groups from Argentina, and the Brazilian group. Isoenzymatic data revealed three diagnostic loci among individuals from Brazil and Argentina, and one locus between both Argentinian groups. At the chromosomal level (amount of autosomal constitutive heterochromatin), both Argentinian groups clearly differed from the Brazilian group, but they were indistinguishable from each other. Considering the preceding studies, Jurberg *et al*. (1998) revalidated once again *T. garciabesi* as a distinct species, involving the individuals from northwest Argentina. This revalidation relied on morphological traits (such as overall size, coloration, head, and genitalia morphology) as well as genetic data.

The variability within *T. sordida* also extended to Bolivia, where two distinct putative species were recognized through multilocus enzyme analyses (Noireau *et al*. 1998). These groups originally identified as Group 1 and Group 2 (now recognized by chromosomal markers as *T. sordida* and *T. garciabesi* respectively) were sympatric in some regions and even interspecific hybrids were identified (Panzera *et al*. 2015). Several years later, quantitative and qualitative analyses of the cuticular hydrocarbons (Calderón-Fernández and Juárez 2013) recognized three different groups within *T. sordida*: *T. garciabesi*, *T. sordida* from Brazil and *T. sordida* from Argentina, similar as described by Panzera *et al*. (1997). In Paraguay, comparative analyses of *T. sordida* populations from western and eastern regions reveal striking differences in the random amplified polymorphic DNA profiles (RAPD), head and wing morphometric and feeding patterns (González-Brítez *et al*. 2014). These authors attribute these differences to population polymorphism caused by eco-geographical isolation by distance.

In 2015, a cytogenetic study involving 139 individuals using chromosomal markers — 45S rDNA location and C-banding— confirmed the presence of four distinct and well-defined groups within *T. sordida sensu lato (s.l.)*: *T. garciabesi* (samples from northwest and central Argentina, western Paraguay and the Bolivian Chaco), *T. sordida sensu stricto (s.s.)* (samples from Brazil, Central and Eastern Paraguay, and the Bolivian Chaco), *T. sordida* Argentina (specimens from northeast Argentina, eastern Paraguay, and the Bolivian high valleys) and *T. sordida* La Paz (domestic individuals from the highlands of La Paz, Bolivia) (Panzera *et al*. 2015). Based also on *cytochrome oxidase subunit 1* (*coI*) sequences data, this study proposed the existence of a new species, widely distributed in the northeast of Argentina (named *T. sordida* Argentina).

Based on individuals from natural populations, morphometric shape measurements for the head, wings and pronotum, differentiated *T. sordida* Argentina from *T. sordida* from Brazil (plus Bolivia), as well as from *T. garciabesi* (Gurgel-Goncalves *et al*. 2011, Nattero *et al*. 2017). However, both studies (independently conducted and using material from different geographical locations), agreed that a high percentage of individuals from different species overlap in their measurements, which could lead to erroneous identification of several individuals using these characters. A similar conclusion was obtained with geometric morphometric of the hemelytra (Belintani *et al*. 2020). According to Nattero *et al*. (2017), this misidentification can be attributed to various factors, including local adaptation or genetic drift, a high level of phenotypic plasticity, morphological convergence, or even natural hybridization.

Finally, in 2020, *T. sordida* from Argentina was formally described as a new species, named *T. rosai*. This taxonomic designation was based on the previously mentioned knowledge and new analyses encompassing morphology, crossbreeding, morphometric, and molecular data (Alevi *et al*. 2020). The morphological characterization of this novel species involved a comparative study of individuals from an Argentine population (Corrientes, San Miguel, designated as the holotype for *T. rosai*) and individuals of *T. sordida s.s.* from Brazil (Minas Gerais).

Although there are morphological keys that allow the identification of *T. sordida*, *T. garciabesi* and *T. rosai*, their practical application still remains as a challenging task. The recognition and differentiation of each of these three species have been and continue to be subject to controversies due to their high morphological similarity, partially overlapping geographical distributions, and even the existence of natural hybrids among them. Molecular identification and evolutionary relationships among these species and with *T. guasayana* have also proven to be highly problematic due to contradictory results. For example, phylogenetic trees with a *coI* gene fragment showed that *T. sordida*, *T. garciabesi* and *T. rosai* form a monophyletic group, with *T. guasayana* as an outgroup (Panzera *et al*. 2015). Similar results were reported using concatenated mitochondrial and nuclear genes: *coI, cytochrome b* (*cytb*), 16S, 18S, and 28S (Belintani *et al*. 2020). However, a study from the same research group that employed nearly identical markers and sequences (*coI*, *cytb*, 16S, 28S, and ITS-2) place *T. guasayana* within the same clade as *T. sordida* and its closely related species (Alevi *et al*. 2020).

All these species also differ in their epidemiological importance as vectors of Chagas disease. *Triatoma sordida* exhibits significant epidemiological variation across its distribution range in terms of infection rates with *T. cruzi* and its ability to colonize domestic and peridomestic habitats. This variation has been documented in different countries, including Brazil (Guilherme *et al*. 2001, Gurgel-Gonçalves *et al*. 2011, Maeda *et al*. 2012, Dantas *et al*. 2018), Bolivia (Noireau *et al*. 1995), and Paraguay (Sánchez *et al*. 2016, Gonzalez-Britez *et al*. 2021). *Triatoma rosai* populations were primarily found in chicken coops, showing a limited ability to colonize and a low rate of infestation in human dwellings (Bar *et al*. 2001, Rodríguez Planes *et al*. 2018). *Triatoma garciabesi* has been found in nests of Furnariidae and Psittacidae birds, with few records in peridomestic environments (Abrahan *et al*. 2016, Cavallo *et al*. 2016).

This study focuses on phylogenetic and population genetic analyses using *cytb* sequences of *T. sordida*, *T. garciabesi*, and *T. rosai*, including not only new sequences from several natural populations from Argentina, Bolivia, Brazil and Paraguay, but also the complete dataset of sequences reported as *T. sordida* and related species deposited in GenBank. We also included *T. guasayana*, a species often mistaken with *T. sordida,* due to their significant morphological resemblance and overlapping geographic ranges (Gorla *et al*. 1993). These *cytb* studies were complemented, whenever possible, with analyses on the same individuals using previously established chromosomal markers that differentiate the aforementioned species. The aim of this study is to elucidate the molecular differentiation among these species and properly establish their geographical distribution ranges. Given the variability in vector competence among these species and the frequent taxonomic confusion, achieving a more accurate geographical distribution for each species could be advantageous in evaluating the transmission risks of Chagas disease associated with each species in the Southern Cone of South America.

## MATERIALS AND METHODS

### Insects and collection sites

A total of 95 male adults from natural populations of 43 localities from Argentina, Bolivia, Brazil and Paraguay (Table S1, Fig. 1) were studied. Some of these samples —25 individuals— were also analyzed by the chromosomal markers detailed in Panzera *et al*. (2015) (depicted in red in Fig. 1). All insects were collected from peridomestic structures, except intradomestic individuals from Inquisivi (La Paz, Bolivia) and sylvatic ones from Reserva Natural y Cultural Bosques Telteca (Mendoza, Argentina). Using the available morphological keys (Lent and Wygodzinsky 1979, Jurberg *et al*. 1998, Alevi *et al*. 2020), all specimens were identified as members of the Sordida subcomplex.

**Figure 1:**
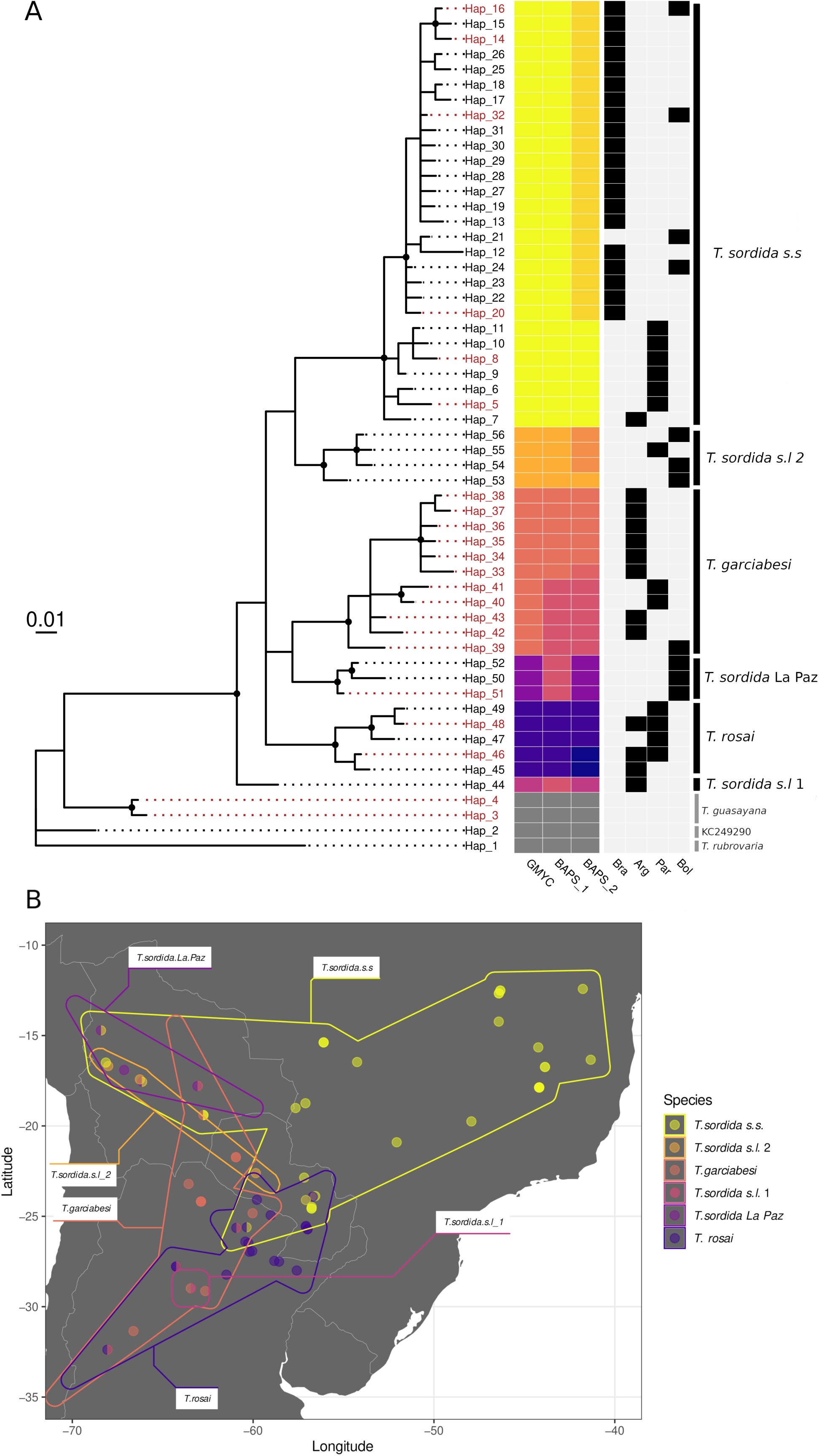
**(A)** Bayesian inference phylogenetic tree obtained from *cytochrome b* fragment (233 bp). Posterior probabilities: support is depicted with black dots over the nodes when above 0.95. In red are the haplotypes that have been cytogenetically analyzed. Colors on the GMYC column define the mitochondrial lineages and used in part B and in figure S3. The two columns on the right represent results from hierBAPS algorithm, different tones of the same colors mean discrepancy with GMYC. The dark gray color represents the outgroup. The country distribution is represented by black squares on the light gray columns. **(B)** Map from the Southern Cone in South America, indicating the geographical distribution of six clades. When two clades are in the same locality, the circle is splitted in two.

### DNA extraction and sequencing

For DNA extraction, three legs preserved in 70% ethanol were used from each specimen and total DNA was extracted by a standard phenol-chloroform procedure. The mitochondrial *cytb* gene was selected for the analyses as it has been proven to be successful for the study of other species complexes within Triatominae (Monteiro et al. 2003), and because it is the mitochondrial fragment with the highest number of sequences available in the GenBank. A fragment of about 500bp of the mitochondrial *cytb* gene was amplified by PCR using primers CYTB7432F (Monteiro *et al*. 2003) and Cob-82R-DEGEN (Patterson and Gaunt 2010). Amplifications were generated by 30/35 cycles of 30s at 95°C, 30s at 58/47°C and 1 min at 72°C, preceded by 5 min at 95°C and followed by 10 min at 72°C. The PCR products were sent to Macrogen Inc. (Korea) for DNA purification and sequencing. Sequences were manually curated by chromatogram evaluation using Chromas (https://technelysium.com.au/wp/chromas/), and then deposited in the GenBank database (http://www.ncbi.nlm.nih.gov), under accession numbers PP972075 to PP972104 (Table S2).

### DNA sequence analyses

Given the high morphological similarity among *T. sordida*, *T. garciabesi*, *T. rosai*, and even these species with *T. guasayana*, we conducted a NCBI search of all GenBank sequences identified as *T. sordida*, *T. garciabesi*, and *T. guasayana*. This strategy allowed us to recognize potentially misidentified bugs, based on morphological characters. Both sequenced individuals of *T. guasayana* (PP972075 and PP972076) were identified as belonging to this species according to the chromosomal markers described by Panzera *et al*. (2015).

We sequenced a fragment of approximately 500 bp. However, in order to include a larger number of the GenBank sequences, genetic analyses were performed using a 233 bp fragment. Several sequences of great importance deposited in GenBank (*i.e.* sequences KR822185 to KR822199) were short, and only included the central region of our dataset. We already experienced the same problem with other species, *Triatoma maculata* (Erichson 1848). And demonstrated that the long and short fragments of the *cytb* gene retrieved the same information (Gomez-Palacio et al 2023). As an outgroup, we selected *T. rubrovaria* (Blanchard, 1843), a species morphologically well differentiated from Sordida subcomplex species but belonging to an evolutionarily close subcomplex —also the same as *T. guasayana*. Sequence alignment was performed using MAFFT v7.310 (Katoh and Standley 2013).

The R core base version 3.6.1 (R core team 2013), and R package *phangorn* (Schliep 2011) were used to estimate a phylogenetic maximum likelihood (ML) tree with 1000 bootstrap pseudoreplicates for node support. Bayesian Inference (BI) was implemented in MrBayes3.2.7 (Ronquist *et al*. 2012). For BI we used four Markov chains for 12 runs of 20 million iterations. The JModeltest function implemented in the *phangorn* package was used to find the best-fitted nucleotide substitution model, and decisions were taken under the Bayesian Information Criterion (BIC). The phylogenetic trees were visualized and edited in the R package *ggtree* (Yu *et al*. 2017). Next, clades within the dataset were defined, employing Hierarchical Bayesian Analysis of Population Structure (HierBAPS) implemented in the R package *rhierbaps* (Cheng *et al*. 2013, Tonkin-Hill *et al*. 2018). In order to have an integrated view of putative species delimitation, the Generalized Mixed Yule Coalescent (GMYC) method (Pons *et al*. 2006) was applied. For this purpose, an ultrametric tree was employed, which was based on a Yule process and obtained using BEASTv.2.3.7 (Bouckaert *et al*. 2014). GMYC was carried out in R using *ape* (Paradis *et al*. 2004) and *splits* packages (Ezard *et al*. 2009). The length of the MCMC chain was 100 million iterations sampling every 10,000. Stationarity and convergence of all parameters was verified using the Tracer v 1.5 software (Rambaut *et al*. 2018).

The R core base version 3.6.1 (R core team 2013), *vegan* (https://CRAN.R-project.org/package=vegan), *ape* (Paradis *et al*. 2004) and *pegas* (Paradis 2010) packages were employed to calculate the within and intergroup pairwise Kimura two parameters (K2p) distance, Tajimas’ D test (Tajima 1989) as well as nucleotide diversity (π), number of haplotypes (h), segregating sites (S) and haplotype diversity (H). Finally, we construct a minimum spanning tree (MST) haplotype network using PopART software (Leigh and Bryant 2015).

### Chromosomal studies

Whenever possible, to complement molecular identification, individuals were also analyzed using chromosomal markers. The chromosomal identification criteria for each species included chromosomal position of ribosomal clusters (by fluorescent *in situ* hybridization) and distribution of C-heterochromatin (by C-banding), as previously described in Panzera *et al*. (2015).

## RESULTS

The total analyzed dataset included 56 haplotypes corresponding to *T. sordida*, *T. garciabesi*, *T. rosai*, *T. guasayana* and *T. rubrovaria* (Fig. 1A, Table 1, Table S2). Among these, 31 haplotypes correspond to this study, while the other 25 haplotypes were sequences extracted from GenBank (Table S2).

**Table 1:** Mitochondrial DNA sequence diversity among the six clades of Sordida subcomplex identified by phylogenetic analyses plus *Triatoma guasayana* as an outgroup. Estimates derived from analysis of *cytochrome b* fragment (233 bp).

| Species | N | Nh | S | Hd | pi | Tajimas' D | Tajimas' D p-value |
| --- | --- | --- | --- | --- | --- | --- | --- |
| *T. sordida sensu stricto* | 160 | 28 | 32 | 0.706 | 0.007 | -2.132 | 0.033 |
| *T. garciabesi* | 22 | 11 | 25 | 0.909 | 0.027 | -0.371 | 0.711 |
| *T. sordida sensu lato 1* | 1 | 1 | 0 | NA | NA | NA | NA |
| *T. rosai* | 48 | 5 | 8 | 0.368 | 0.007 | -0.167 | 0.867 |
| *T. sordida* La Paz | 4 | 3 | 4 | 0.833 | 0.009 | -0.780 | 0.435 |
| *T. sordida sensu lato 2* | 4 | 4 | 7 | 1.000 | 0.015 | -0.817 | 0.414 |
| *T. guasayana* | 2 | 2 | 1 | 1.000 | 0.004 | NA | NA |

Note: N: number of specimens analyzed; Nh: number of haplotypes; S: number of segregating sites; Hd: haplotype diversity; $\pi$ : nucleotide diversity; Tajimas' D neutrality test value and its p-value.

GMYC and hierBAPS algorithms retrieved similar results from each other. Differences between GMYC and hierBAPS level 1 involved *T. garciabesi, T. sordida* La Paz and *T*. *sordida sensu lato* 1 (*T. sordida s.l.* 1). These are resolved if hierBAPS level 2 is considered. The exception is that *T. garciabesi* is split into two clades. This could be reflecting the division between both lineages within *T. garciabesi* described recently (Verly *et al*. 2024). Unfortunately, we have not enough samples to test this hypothesis in the present paper. Therefore, GMYC results were taken as the best option to describe the phylogenetic groups or clades.

According to our analysis, seven GenBank sequences were misidentified (Table S1, in red). Surprisingly, a *T. sordida* GenBank sequence (Hap. 1 —KC249290) was also included within the outgroup, presenting a high identity (98.71%) with *T. infestans* (Klug, 1834) (Acc. Number HQ333230). Two *T. sordida* GenBank sequences (KC249293 and KC249295) were identified as *T. garciabesi* and *T. rosai*, respectively. Finally, other four GenBank sequences erroneously identified as belonging to *T. guasayana* grouped into two clades within the Sordida subcomplex (*T. sordida s.s.* and *T. sordida* La Paz (Fig. 1A y Fig. S1).

According to the topology of the BI (Fig. 1A) and ML trees (Fig. 1S) we recognized six clades within the Sordida subcomplex. In the ML tree all clades form a basal polytomy, hence phylogenetic relationships at the base of the tree were not resolved (Fig. 1S). In the BI tree, a similar basal polytomy is found after the divergence of the Hap. 44, which belongs to *T. sordida s.l.* 1 (Fig. 1A).

In Figure 1, the first clade from top to bottom (depicted in yellow —haplotypes 5 to 32) corresponds to *T. sordida s.s.* and is located mainly in Brazil, but also in Bolivia, Paraguay and one haplotype (Hap. 7) in Argentina (detailed also in Fig. S2). The second clade (in pale orange —haplotypes 53 to 56), corresponds to *T. sordida sensu lato* 2 *(T. sordida s.l.* 2), distributed in Bolivia and Paraguay. The third clade (colored in dark orange —haplotypes 33 to 43), is *T*. *garciabesi*, located mainly in Argentina but also in Paraguay and Bolivia. Fourth clade (colored in light purple, haplotypes 50 to 52), corresponds to *T*. *sordida* La Paz, which is distributed exclusively in Bolivia. The fifth clade, (dark purple —haplotypes 45 to 49), is *T. rosai* (*T. sordida* Argentina *sensu* Panzera *et al*. 2015), which is distributed mainly in Argentina, but also in Paraguay. The last clade at the bottom of the in-group (seance color) possesses only haplotype 44 located in Argentina, and receives the name of *T. sordida s.l.* 1. In Supplementary Figure 2, the detailed geographical distribution of each haplotype is shown, discriminated by clades with the same colors as in Figure 1.

All clades retrieved by GMYC and hierBAPS algorithms are clearly represented in the minimum spanning tree (MST) network (Fig. S3). It could be seen how mutational steps are larger in the branches that separate each group. It is important to note that the haplotype 44 (a singleton of *T. sordida s.l.* 1) is separated from the other haplotypes by several mutational steps (Fig. S3). The star-like shape of the haplotype network for *T. sordida s.s.* is also particular, featuring a central variant encircled by haplotypes exhibiting only minor mutational differences compared to the structure observed in other species such as *T. garciabesi*.

The greatest number of haplotypes (28) and segregating sites (32) was found in *T. sordida s.s.* (Table 1). However, the highest haplotype diversity was in *T. garciabesi* (0.91) and *T. sordida s.l.* 2 (1.0) and the lowest in *T. rosai* (0.37). The highest nucleotide diversity value was 0.027 in *T. garciabesi,* and the lowest 0.007 in *T. sordida s.s.* and *T. sordida* La Paz. Tajimas’ D was significant only for *T. sordida s.s.* (-2.132, p = 0.03) (Table 1).

According to K2p corrected genetic distances (Table 2), a *T. sordida* GenBank sequence (Hap. 1 —KC249290) displays genetic distances higher than 21% with any of the Sordida subcomplex species. The smallest genetic distance observed for haplotype 1 (erroneously identified as *T. sordida*, likely *T. infestans*) was 16% with *T. rubrovaria*. Regarding the two-outgroup species, *T. rubrovaria* and *T. guasayana*, they display genetic distances ranging from 12% to 19% when compared to Sordida subcomplex species. Within the Sordida subcomplex, the greatest distance was between *T. sordida s.s.* and *T. garciabesi* (10.6%), and the smallest was 6.1% (*T. sordida s.l.* 2 with *T. sordida* La Paz and *T. sordida s.l.* 1; and *T. sordida s.l.* 1 versus *T. rosai* and *T. sordida* La Paz). The intra-clade variation ranges between 1.2% and 3.3%, as seen in *T. sordida* La Paz and *T. garciabesi,* respectively (Table 2). Genetic distances results are also given for each haplotype on Table SII.

**Table 2.** Mean Kimura 2-parameter (K2p) distances (expressed in percentages) of *cytb* sequences among the clades of Figure 1, plus *Triatoma sordida* GenBank (Hap. 1 - KC249290), *T. rubrovaria* and *T. guasayana* as outgroups. Mean within-clade distances are in bold on the diagonal.

|  | Hap. 1<br>KC24929<br>0 | *T.*<br>*rubr.* | *T.*<br>*gua.* | *T. gar.* | *T.*<br>*ros.* | *T.*<br>*sor.*<br>*La*<br>*Paz* | *T. sor.*<br>*s.l. 1* | *T. sor.*<br>*s.l. 2* | *T.*<br>*sor.*<br>*s.s..* |
| --- | --- | --- | --- | --- | --- | --- | --- | --- | --- |
| *T. rubrovaria* | 15.99 | **NA** |  |  |  |  |  |  |  |
| *T. guasayana* | 17.39 | 6.63 | **0.43** |  |  |  |  |  |  |
| *T. garciabesi* | 23.35 | 13.88 | 13.35 | **3.25** |  |  |  |  |  |
| *T. rosai* | 23.14 | 15.04 | 16.54 | 10.09 | **2.02** |  |  |  |  |
| *T. sordida* La | 22.43 | 14.07 | 12.26 | 7.09 | 7.60 | **1.16** |  |  |  |
| *T. sordida sensu*<br>*lato 1* | 21.07 | 12.75 | 12.49 | 6.97 | 6.34 | 6.05 | **NA** |  |  |
| *T. sordida sensu*<br>*lato 2* | 22.87 | 13.91 | 14.99 | 7.82 | 7.16 | 6.11 | 6.50 | **1.54** |  |
| *T. sordida sensu*<br>*stricto* | 24.62 | 14.58 | 18.99 | 10.61 | 8.37 | 9.91 | 9.44 | 8.34 | **2.04** |

Chromosome analyses showed no novel patterns. All analyzed individuals presented the same chromosomal characteristics described in Panzera *et al*. (2015). Here, we obtained new results for *T. sordida s.s, T. garciabesi* and *T. rosai* individuals. For *T. sordida s.s*: autosomal C-heterochromatin in several autosomal pairs and 45S ribosomal clusters on the X chromosome. *Triatoma garciabesi*: no autosomal C-heterochromatin and ribosomal clusters on the X chromosome. *Triatoma rosai*: no autosomal C-heterochromatin and 45S ribosomal clusters present on both XY sex chromosomes.

## DISCUSSION

Based on a broad geographic sampling, this investigation report evidence of an extensive genetic diversity involving *T. sordida* and related species, highlighting the following conclusions:

### Phylogenetic groupings

Within the Sordida subcomplex species, we recognize six clades with pairwise *cytb* K2p distances greater than 6%, and with support values (BPP) higher than 0.90 (Table 1, Fig. S1). The three clades are formally recognized as true species (*T. sordida s.s.*, *T. garciabesi* and *T. rosai*) showed genetic distances between 8.4% to 10.6 %, very similar as those reported by Alevi *et al*. (2020) (7.4% to 9.7%). The other three clades, *T. sordida s.l.* 1, *T. sordida s.l.* 2, and *T. sordida* La Paz, exhibit genetic distances between 6 to 11%. Studies of DNA barcoding and DNA taxonomy have proposed that threshold values of sequence divergence can assist in delineating animal species (Hebert *et al*. 2003). Anyway, the divergence thresholds are arbitrary and can vary widely depending on the genetic marker employed and even among different insect groups (Meier *et al*. 2006). Mitogenome analyses of 90 hemipteran species indicated that *cytb* and *coI* gene fragments undergo evolution at comparable rates (Wang *et al*. 2015). However, within triatomines, *cytb* gene evolves slightly more rapidly than *coI* (Pfeiler *et al*. 2006, Pita *et al*. 2021). In *Triatoma* sister species, *cytb* sequence divergence levels (measured by K2p distances) generally exceed 7.5% (Monteiro *et al*. 2004, 2013, Pfeiler *et al*. 2006), significantly surpassing the 2–3% threshold commonly used in several insect groups, including true bugs (Meier *et al*. 2006, Raupach *et al*. 2014). In our analyses, *cytb* K2p distances range from 6% to 11% among the six clades here identified, whereas within-clades distances do not exceed 3.3% (Table 2). Although the three clades not recognized as valid species fall just below the 7.5% threshold (*T. sordida* La Paz, *T. sordida s.l.* 1, and *T. sordida s.l.* 2), these results suggest strong genetic differentiation within the species now recognized as *T. sordida.* If we compare these three taxa exclusively with *T. sordida s.s.*, the genetic distances are always greater than 8.3% (range 8.3% to 9.9%). More extensive population analyses with various nuclear and mitochondrial markers could clarify whether these three clades involve new species or not.

### Incorrect Molecular Species Identification

This work highlights the incorrect taxonomic identification of numerous *cytb* sequences from GenBank, as it has been reported for other mitochondrial DNA sequences (Panzera *et al*. 2015, Belintani *et al*. 2020) (Fig. 1 and Table 3S). The most striking example is the haplotype 1 identified as *T. sordida* (KC249290), with such a high level of molecular divergence that it even appears as an outgroup in our analyses (Figure 1 and Table 2). Regarding *T. guasayana,* the four GenBank sequences named would not belong to this species. Furthermore, *T. guasayana* from GenBank were not clustered with individuals previously identified as *T. guasayana* based on chromosomal characteristics reported by Panzera *et al*. (2015) —absence of autosomal heterochromatin and ribosomal DNA clusters on a single autosomal pair. These incorrect identifications on GenBank accessions explain the contradictory results of numerous reports whether *T. guasayana* is within or outside the Sordida subcomplex (Justi *et al*. 2014, Alevi *et al*. 2020) above commented in the “Introduction” section.

### Geographical distribution and genetic variability within each clade

***T. sordida sensu stricto*:** This species is widely distributed in Brazil, Paraguay, Bolivia (highlands and Chaco region). One specimen from Argentina (Chaco, El Colchon, Hap. 7) (Fig. 1) is described but could be a case of introgression. Furthermore, it is sympatric with the haplotype 48 of *T. rosai. Triatoma sordida s.s.* also possess overlapped distribution with clades *T. garciabesi*, *T. sordida* La Paz, and *T. sordida s.l.* 2 in Bolivia. All the sordida-like specimens found in Brazil belong to this clade (Fig. S2A). Several haplotypes were confirmed by chromosomal markers (identified in red in Fig. 1). Despite its widespread geographic distribution and the large number of individuals analyzed, along with a high number of haplotypes and segregating sites (Table 1), this species exhibits the lowest nucleotide diversity among the six identified clades, with K2p interclade distances at 2.0% (Table 2). This narrow molecular variability aligns with previous isoenzymatic and DNA studies (Monteiro *et al*. 2009, Pessoa *et al*. 2016, Madeira *et al*. 2020). The K2p genetic distances of *T. sordida s.s.* with the remaining five clades are greater than 8.3%, including the three taxa not formally recognized as species (Table 2). A significant negative Tajima’s D value and the star-shaped haplotype network for this species suggest a recent process of population expansion, indicative of a strong colonizing capability in this taxon relative to others within the subcomplex. The same individuals previously identified as chromosomal hybrids by Panzera *et al*. (2015) from Izozog (Bolivia) were examined and found to have *T. sordida s.s. cytb* haplotypes (Hap. 16 and 32).

***T. sordida sensu lato* 2:** The clade is distributed in Paraguay (Boquerón Department) (haplotype 55), and three GenBank sequences from highlands of Bolivia (Hap. 53, 54 and 56) (Fig. S2B). In Bolivia, this taxon occurs in the same geographic areas as *T. sordida s.s.* and *T. sordida* La Paz, from which it is separated by K2p distances of 8.3% and 6.1%, respectively. Chromosomal analyses were not possible to perform for this taxon, as their gonads were not collected while the insects were still alive.

***T. sordida sensu lato* 1:** The clade is restricted to a single sequence (Hap. 44), separated for the other haplotypes by several mutational steps (Fig. S3), from an individual localized in the humid Chaco region from Argentina (Santiago del Estero, Salavina) (Fig. S2D). Other individuals from the Sordida subcomplex that are geographically closest are identified as *T. garciabesi*, with genetic distances approximately 7% (Table 2). It is noteworthy that this individual shows a considerable genetic distance from *T. sordida s.s.* (9.44%), indicating that it clearly represents a different taxon. *Triatoma sordida s.l.*1 is also distant to *T. sordida* La Paz and *T. sordida s.l.* 2 (K2p distances of 6.1% and 6.5%, respectively), which are geographically very distant (Fig. S2). Chromosomal analyses were not possible to carry out for this taxon either.

***T. garciabesi:*** This species is common and widely distributed in northern Argentina, although it is also found in a locality in Paraguay (Boquerón, haplotypes 40 and 41) and in the Bolivian Chaco (Hap. 39) (Fig. 1 and Fig. S2D). In several localities of Argentina, this species is found in the same areas as *T. rosai*. This clade is characterized by presenting the highest nucleotide and haplotype diversity among the 6 clades analyzed (Table 1), with an intraclade K2p distance of 3.3% (Table 2). This high intraspecific variation with *cytb* is very similar to that recently described involving *coI* fragments (Verly *et al*. 2024). The eleven haplotypes identified in this species were chromosomally analyzed (Fig. 1).

***T. rosai:*** This study detects this species in several provinces of Argentina (some of which were previously undescribed) and also in several departments of Paraguay (Fig. 1; Fig. S2E; Table S1). Chromosomal and cuticular hydrocarbon profiles have identified the presence of this taxon also in Bolivia (Cochabamba and Santa Cruz Departments) (Panzera *et al*. 2015, Moriconi *et al*. 2022). Despite this wide distribution, the species exhibits a low number of haplotypes with the least nucleotide diversity (Table 1). Several individuals of this species (haplotypes 46 and 48) were examined, and corroborated via chromosomal markers (Fig. 1A).

***T. sordida* La Paz:** This taxon is found exclusively in Bolivia (La Paz, Cochabamba and Santa Cruz) and comprises three haplotypes (50, 51, and 52) (Fig. 1, Fig. S2E). Two of them are extracted from GenBank and classified as *T. guasayana.* The third one, Hap. 51, described here from an individual that was chromosomally analyzed in Panzera *et al*. (2015), presents 45S ribosomal clusters in an autosomal pair and C-heterochromatic autosomes, distinctive traits of *T. sordida* La Paz. Our analysis places these three haplotypes within the same clade, which is genetically quite distinct from *T. guasayana* (K2p = 12.3%) and *T. sordida s.s.* (9.9%).

A recent research proposed that *T. sordida* La Paz was not a valid species due it low genetic distance with *T. sordida s.s.* (Madeira *et al*. 2021). To verify this conclusion, the same *cytb* sequences from Madeira *et al*. (2021) were used here and confirmed that these individuals of *T. sordida* from the La Paz region are in fact *T. sordida s.s.* (haplotype 32 = MZ700100 and haplotype 21 = MZ700101). Madeira *et al*. (2021) failed to assume that all *T. sordida* individuals from La Paz region belonged to the lineage *T. sordida* La Paz. Our molecular and chromosomal data indicate the coexistence of both clades, *T. sordida s.s.* and *T. sordida* La Paz, in the La Paz region, with significant genetic distinctions between them (K2p distances of approximately 10%). This substantiates the need for an exhaustive morphological analysis of *T. sordida* La Paz, recognizing it as a separate species. Cuticular hydrocarbons studies on individuals from Cochabamba also suggest the existence of other species beyond those already described (Moriconi *et al*. 2022), possibly *T. sordida* La Paz or *T. sordida s.l.* 2, as proposed here.

## CONCLUSION

Building on the analysis of a mitochondrial *cytb* gene fragment supplemented by chromosomal analyses, our study identifies six molecular taxa within the Sordida subcomplex. Three of these taxa are formally recognized as species (*T. sordida s.s.*, *T. garciabesi*, and *T. rosai*), while the others (*T. sordida s.l. 1*, *T. sordida s.l. 2*, and *T. sordida La Paz*) require further investigation to confirm their taxonomic status. Notably, four of these clades also exhibit chromosomal differences previously described by Panzera et al. (2015), underscoring the complexity of this group.

Our study not only provides a detailed molecular and chromosomal framework for understanding the Sordida subcomplex but also emphasizes the critical role of integrating multidisciplinary approaches to resolve taxonomic ambiguities. The accurate identification of species and clades within this complex is essential for understanding their evolutionary trajectories and their role in the transmission dynamics of *Trypanosoma cruzi*. Misidentifications in public databases, such as GenBank, and the existence of natural hybrids further complicate phylogenetic analyses, highlighting the necessity of robust taxonomic tools in vector control programs.

Addressing these challenges requires significant collaboration among research groups, leveraging advanced genomic tools such as SNP analyses and comprehensive chromosomal studies. This approach could not only enhance our understanding of genetic flow and hybridization but also serve as a model for resolving taxonomic ambiguities in other insect vectors. Moreover, these efforts must overcome practical barriers, including the extensive geographic overlap of clades in regions like the Andean and Gran Chaco areas, and the humid Chaco regions of Paraguay and Argentina, where the collection of samples is logistically and financially demanding.

Our findings also raise intriguing questions about the evolutionary forces driving diversification within the Sordida subcomplex. Future studies could explore whether environmental gradients, host preferences, or historical biogeographic events contribute to the genetic differentiation observed among clades. Conducting comparative analyses of mitochondrial and nuclear markers, alongside chromosomal and SNP data, will be essential to further elucidate the taxonomy of this complex.

## Acknowledgements

We thank E.B. Oscherov, A. Gonzalez and R. Cardozo for providing biological materials used in this study. GBR, NR, FP and SP acknowledge financial support by Programa Desarrollo de Ciencias Básicas (PEDECIBA, Uruguay) and Sistema Nacional de Investigadores (SNI-ANII). JN, RVP, ALCF, PAL and MJC are members of the CONICET Researcher’s Career.

## AUTHORS’ CONTRIBUTIONS

JN, FP and SP **conceived and designed the study**; CML, PAL, ARA, MJC, CSR, MCVG, ALCF and MR **collected bugs in the field**. GBR, NR, BSR and SP performed laboratory work and *in silico* analyses. GBR, JN, NR, RVP, ALCF, PL, FP and SP **drafted the manuscript**. PL, FP and SP **funding acquisition**. All authors read and approved the final version of the manuscript.

## Supplementary Data

**Figure S1:** Maximum likelihood tree obtained from a *cytochrome b* fragment. Values of Bayesian posterior probabilities are depicted with numbers and circles on the nodes.

**Figure S2:** Map of the Southern Cone in South America, illustrating the distribution of each clade identified through phylogenetic analyses. **(A)** *Triatoma sordida sensu stricto* **(B)** *T. sordida sensu lato* 2 **(C)** *T. garciabesi* **(D)** *T. sordida sensu lato* 1 **(E)** *T. sordida* La Paz **(F)** *T. rosai*.

**Figure S3.** Minimum-spanning haplotype network based on a 233 bp *cytb* alignment. The nodes are haplotypes, with node size proportional to haplotype frequency. The numbers of mutational steps separating haplotypes are represented by dashes along the edges.

**Table S1:** Geographic origin of individuals, depicting the mitochondrial haplotype, latitude and longitude, locality and country (when the coordinates used in the map were not the exact ones, are specifically mentioned in the table). Species as are indicated in NCBI and the reference paper. In red appear the GenBank sequences erroneously identified.

**Table S2:** Mitochondrial haplotypes (*cytb*) and GenBank accession numbers of the sequences here analyzed.

**Table S3:** Kimura 2-parameter (K2-p) genetic distances computed by haplotypes.

## DATA AVAILABILITY

All sequences used in this article were deposited in GenBank Nucleotide Database (https://www.ncbi.nlm.nih.gov/genbank/) with accession numbers from PP972075 to PP972104.

