## Supplementary figures and images for "Reconstructing Sordida subcomplex (Hemiptera, Reduviidae, Triatominae) phylogeny across species distribution range"

### Supplementary_fig_1

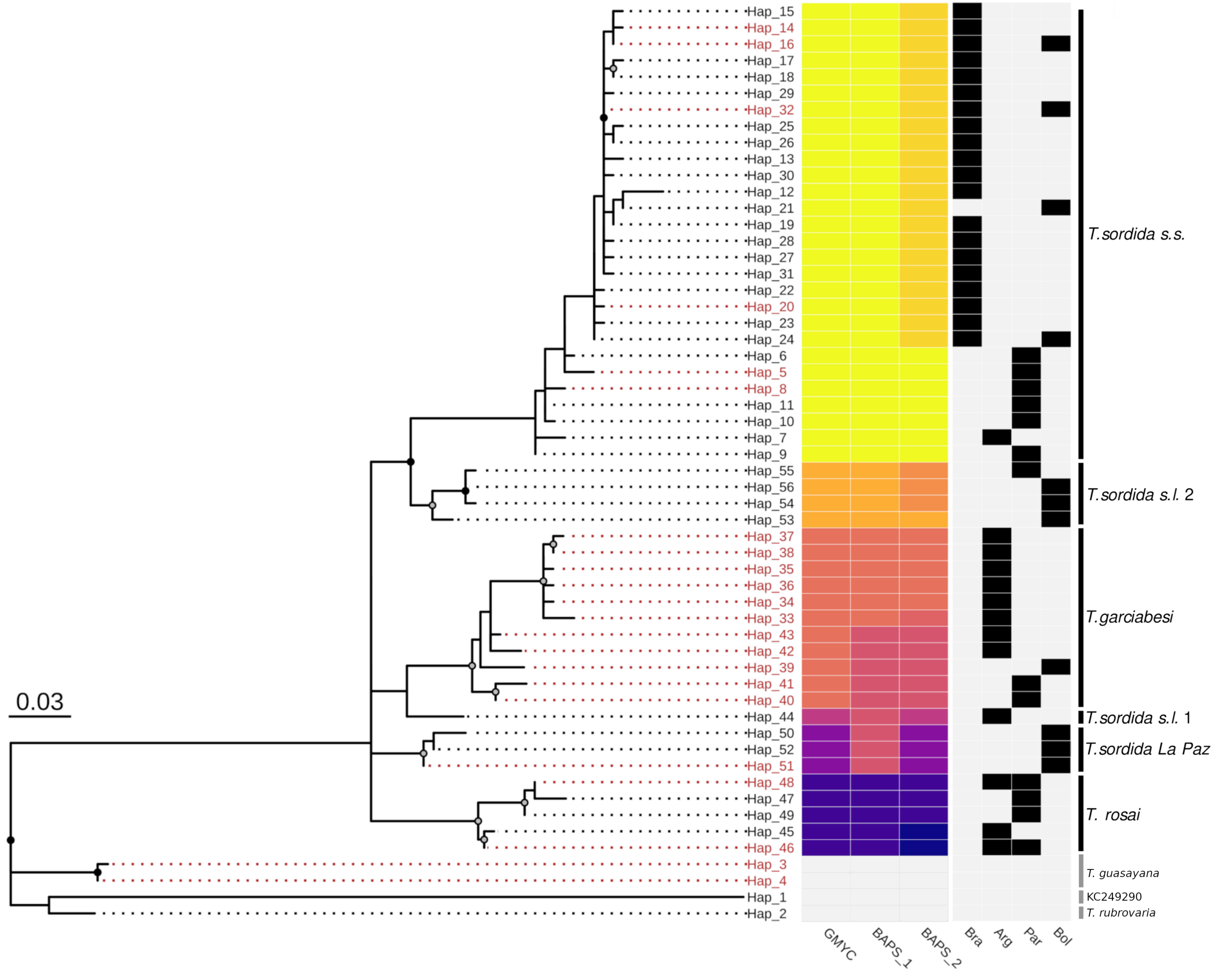

### Supplementary_fig_2

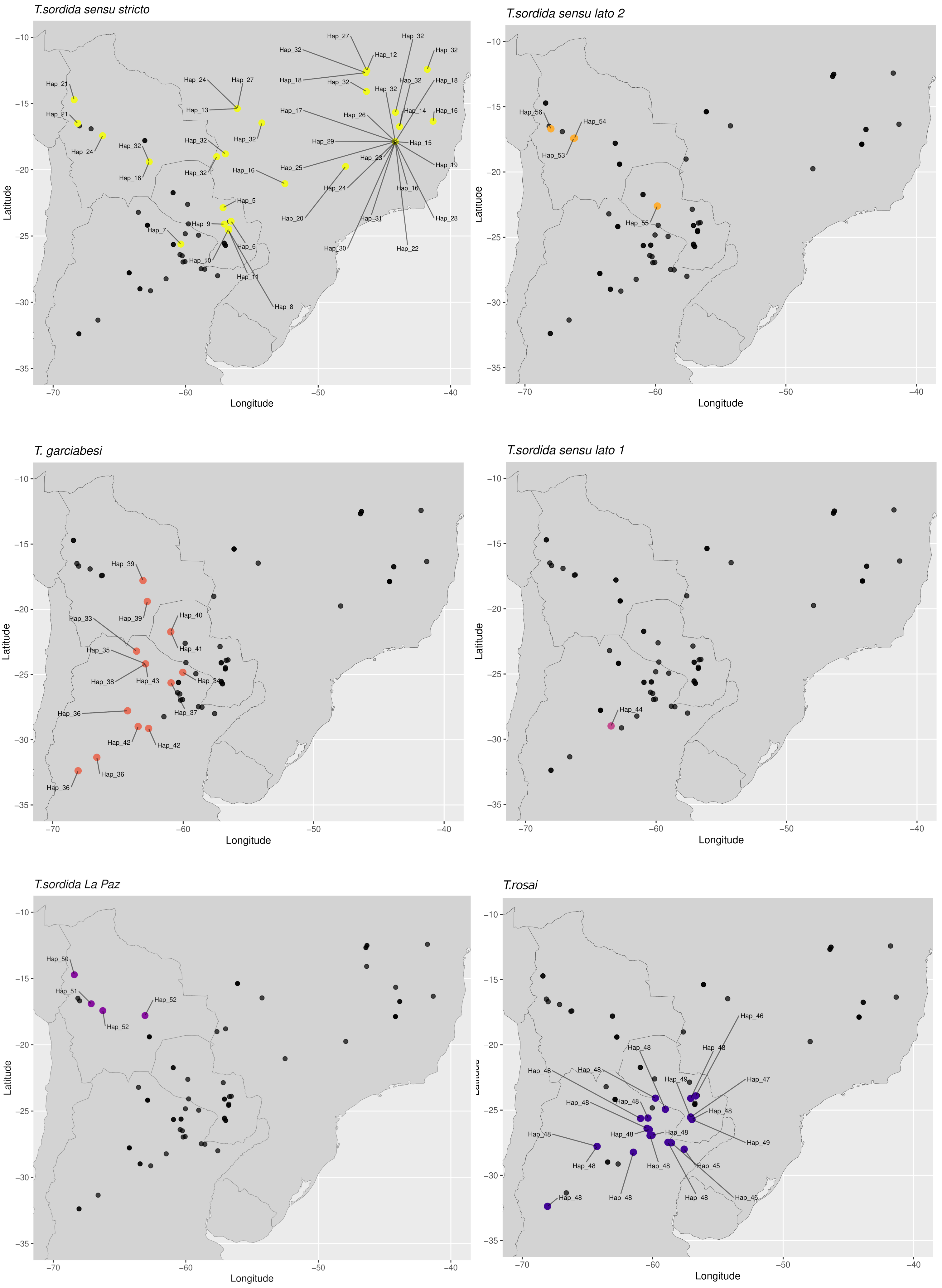

### Supplementary_fig_3

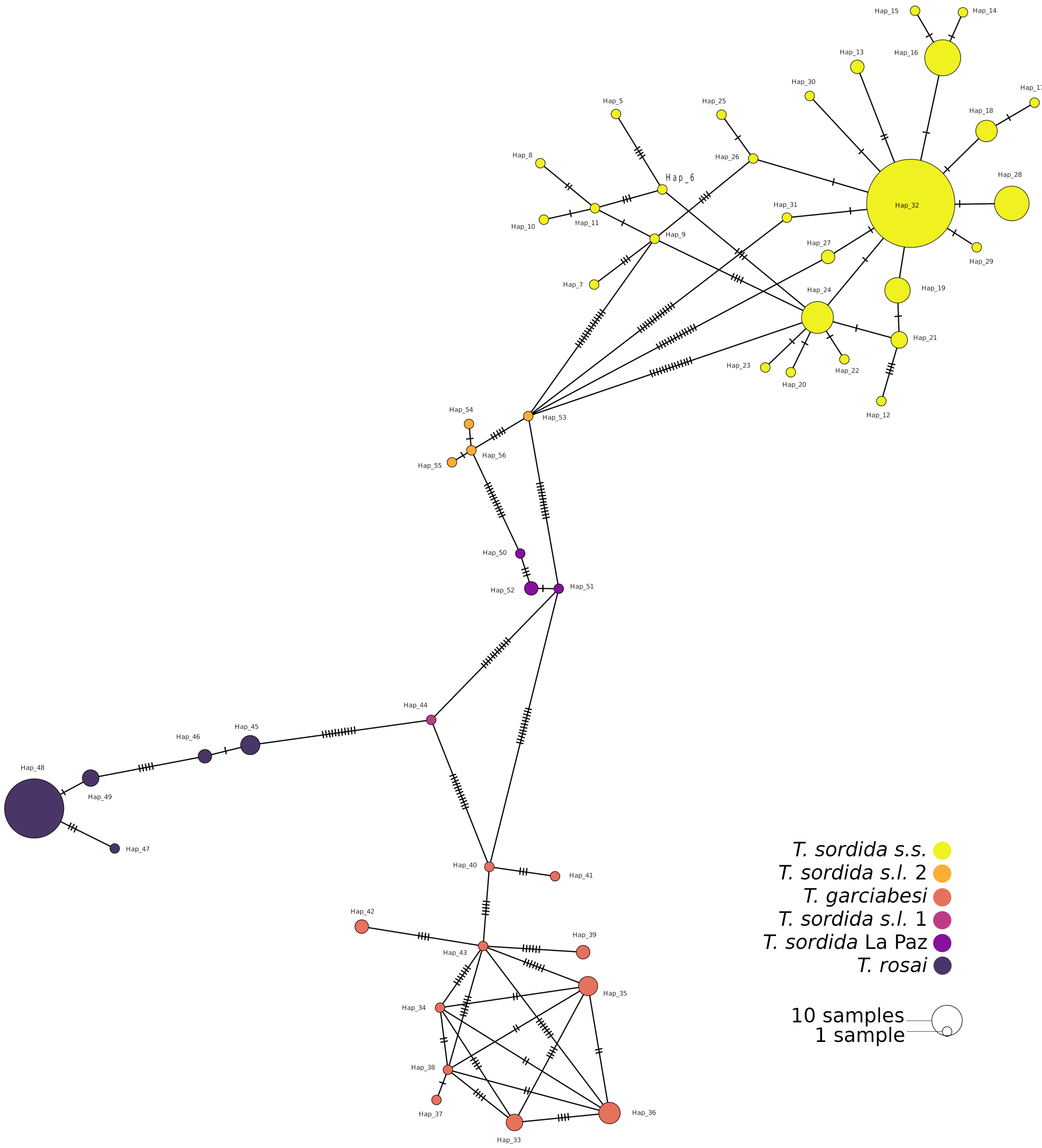
