## Supplementary_table_1 for "Reconstructing Sordida subcomplex (Hemiptera, Reduviidae, Triatominae) phylogeny across species distribution range"

| Haplotype | Lineage | Latitude | Longitude | Geographic origin | NCBI species | Reference |
|---|---|---|---|---|---|---|
| Hap_1 | T. infestans | Not on the map | | | T. sordida | Justi et al. 2014 |
| Hap_2 | T. rubrovaria | Not on the map | | | T. rubrovaria | Justi et al. 2014 |
| Hap_3 | T. guasayana | Not on the map | | Argentina, Córdoba, Sobremonte. | | This paper |
| Hap_4 | T. guasayana | Not on the map | | Argentina, Córdoba, Sobremonte. | | This paper |
| Hap_5 | T.sordida.s.s | -22,86108 | -57,175868 | Paraguay, Concepcion, Zona 3 | | This paper |
| Hap_6 | T.sordida.s.s | -23,8833 | -56,5667 | Paraguay, San Pedro, Santa Rosa, Centro | | This paper |
| Hap_7 | T.sordida.s.s | -26 | -60 | Argentina, Chaco, El Colchon | | This paper |
| Hap_8 | T.sordida.s.s | -24,568524 | -56,769182 | Paraguay, San Pedro, Itacurubi del Rosario, Rios Rugua | | This paper |
| Hap_9 | T.sordida.s.s | -24 | -57 | Paraguay, San Pedro, San Pedro, Yatebu Rugua | | This paper |
| Hap_10 | T.sordida.s.s | -24 | -57 | Paraguay, San Pedro, Itacurubí del Rosario, Campo Virgen | | This paper |
| Hap_11 | T.sordida.s.s | -25 | -57 | Paraguay, San Pedro, Itacurubi del Rosario, Rios Rugua | | This paper |
| Hap_12 | T.sordida.s.s | -12,51 | -46,33 | Brazil, Tocantins, Combinado | | This paper |
| Hap_13 | T.sordida.s.s | -15 | -56 | Brazil, Mato Grosso, Varzea Grande | | This paper |
| Hap_14 | T.sordida.s.s | -16,74 | -43,86 | Brazil, Minas Gerais, Montes Claros | | This paper |
| Hap_15 | T.sordida.s.s | -18 | -44 | Brazil, Minas Gerais, Buenopolis (Not exact locality) | T. sordida | Pessoa et al. 2016 |
| Hap_16 | T.sordida.s.s | -19 | -63 | Bolivia, Santa Cruz, Izozog | | This paper |
| Hap_16 | T.sordida.s.s | -16 | -41 | Brazil, Minas Gerais, Itaobim | | This paper |
| Hap_16 | T.sordida.s.s | -17,87277778 | -44,18 | Brazil, Minas Gerais, Buenopolis (Not exact locality) | T. sordida | Pessoa et al. 2016 |
| Hap_16 | T.sordida.s.s | -21 | -53 | Brazil, Mato Grosso do Sul, Brasilandia (Not exact coordinates) | T. sordida | Gardim et al. 2013 |
| Hap_17 | T.sordida.s.s | -18 | -44 | Brazil, Minas Gerais, Buenopolis (Not exact locality) | T. sordida | Pessoa et al. 2016 |
| Hap_18 | T.sordida.s.s | -18 | -44 | Brazil, Minas Gerais, Buenopolis (Not exact locality) | T. sordida | Pessoa et al. 2016 |
| Hap_18 | T.sordida.s.s | -13 | -46 | Brazil, Tocantins, Aurora de Tocantins | | This paper |
| Hap_19 | T.sordida.s.s | -17,87277778 | -44,18 | Brazil, Minas Gerais, Buenopolis (Not exact locality) | T. sordida | Pessoa et al. 2016 |
| Hap_20 | T.sordida.s.s | -19,75 | -47,93 | Brazil, Minas Gerais. Uberaba | | This paper |
| Hap_21 | T.sordida.s.s | -14,717401 | -68,413711 | Bolivia, La Paz, Apolo | T. sordida | **Unpublished Belintani** |
| Hap_21 | T.sordida.s.s | -16,49416 | -68,147 | Bolivia, La Paz. Colony | T. sordida | Justi et al. 2014/Madeira et al. 2021 |
| Hap_22 | T.sordida.s.s | -17,87277778 | -44,18 | Brazil, Minas Gerais, Buenopolis (Not exact locality) | T. sordida | Pessoa et al. 2016 |
| Hap_23 | T.sordida.s.s | -18 | -44 | Brazil, Minas Gerais, Buenopolis (Not exact locality) | T. sordida | Pessoa et al. 2016 |
| Hap_24 | T.sordida.s.s | -15,38 | -56,1 | Brazil, Mato Grosso, Varzea Grande | | This paper |
| Hap_24 | T.sordida.s.s | -18 | -44 | Brazil, Minas Gerais, Buenopolis (Not exact locality) | T. sordida | Pessoa et al. 2016 |
| Hap_24 | T.sordida.s.s | -17 | -66 | Bolivia, Cochabamba, chaco Tita. Colony. (Not exact coordinates | T. guasayana | Justi et al. 2014 |
| Hap_25 | T.sordida.s.s | -17,87277778 | -44,18 | Brazil, Minas Gerais, Buenopolis (Not exact locality) | T. sordida | Pessoa et al. 2016 |
| Hap_26 | T.sordida.s.s | -18 | -44 | Brazil, Minas Gerais, Buenopolis (Not exact locality) | T. sordida | Pessoa et al. 2016 |
| Hap_27 | T.sordida.s.s | -12,51 | -46,33 | Brazil, Tocantins, Combinado | | This paper |
| Hap_27 | T.sordida.s.s | -15,38 | -56 | Brazil, Mato Grosso, Varzea Grande | | This paper |
| Hap_28 | T.sordida.s.s | -18 | -44 | Brazil, Minas Gerais, Buenopolis (Not exact locality) | T. sordida | Pessoa et al. 2016 |
| Hap_29 | T.sordida.s.s | -18 | -44 | Brazil, Minas Gerais, Buenopolis (Not exact locality) | T. sordida | Pessoa et al. 2016 |
| Hap_30 | T.sordida.s.s | -17,87277778 | -44,18 | Brazil, Minas Gerais, Buenopolis (Not exact locality) | T. sordida | Pessoa et al. 2016 |
| Hap_31 | T.sordida.s.s | -18 | -44 | Brazil, Minas Gerais, Buenopolis (Not exact locality) | T. sordida | Pessoa et al. 2016 |
| Hap_32 | T.sordida.s.s | -19,401315 | -62,750258 | Bolivia, Santa Cruz, Izozog. | | This paper |
| Hap_32 | T.sordida.s.s | -13 | -46 | Brazil, Tocantins, Aurora de Tocantins | | This paper |
| Hap_32 | T.sordida.s.s | -16,74 | -43,86 | Brazil, Minas Gerais, Montes Claros | | This paper |
| Hap_32 | T.sordida.s.s | -16,464419 | -54,249643 | Brazil, Mato Grosso, Sao Jose do Povo | | This paper |
| Hap_32 | T.sordida.s.s | -18 | -44 | Brazil, Minas Gerais, Buenopolis (Not exact locality) | T. sordida | Pessoa et al. 2016 |
| Hap_32 | T.sordida.s.s | -19 | -58 | Brazil, Mato Grosso do Sul, Corumba (Not an exact coordinates) | T. sordida | Madeira et al. 2021 |
| Hap_32 | T.sordida.s.s | -12 | -42 | Brazil, Bahia, Seabra (Not an exact coordinates) | T. sordida | **Unpublished Belintani** |
| Hap_32 | T.sordida.s.s | | | Brazil, Minas Gerais, Monte Azul. Colony. | T. sordida | **Unpublished Belintani** |
| Hap_32 | T.sordida.s.s | -15,663889 | -44,163889 | Brazil, Minas Gerais, Ibiracatu. Colony. | T. sordida | Justi et al. 2014 |
| Hap_32 | T.sordida.s.s | -19 | -57 | Brazil, Mato Grosso do Sul, Pantanal. Colony. ( | T. sordida | Justi et al. 2014 |
| Hap_32 | T.sordida.s.s | -14 | -46 | Brazil, Goias, Posse. Colony. | T. sordida | Justi et al. 2014 |
| Hap_33 | T.garciabesi | -23 | -64 | Argentina. Salta. Hickman | | This paper |
| Hap_34 | T.garciabesi | -24,83 | -60,03 | Argentina, Formosa, Patiño. | | This paper |
| Hap_35 | T.garciabesi | -24,1833333 | -62,88333 | Argentina, Salta, Rivadavia. Colony. (Not exact coordinates) | T. garciabesi | Justi et al. 2014/Unpublished Belintani |
| Hap_35 | T.garciabesi | -24,1833333 | -62,88333 | Argentina, Salta (Not exact locality) | | This paper |
| Hap_36 | T.garciabesi | -31,35 | -66,62 | Argentina, La Rioja, Rosario Vera Peñaloza | | This paper |
| Hap_36 | T.garciabesi | -28 | -64 | Argentina, Santiago del Estero (Not exact locality) | | This paper |
| Hap_36 | T.garciabesi | -32,38138 | -68,05531 | Argentina, Mendoza, Reserva Natural Bosques Telteca | | This paper |
| Hap_37 | T.garciabesi | -26 | -61 | Argentina, Salta, Balbuena | | This paper |
| Hap_38 | T.garciabesi | -24 | -63 | Argentina, Salta, Rivadavia | | This paper |
| Hap_39 | T.garciabesi | -19 | -62,750258 | Bolivia, Santa Cruz, Izozog | | This paper |
| Hap_39 | T.garciabesi | -17,796675 | -63,080339 | Bolivia, Santa Cruz. Colony. | T. sordida | Justi et al. 2014 |
| Hap_40 | T.garciabesi | -22 | -60,95 | Paraguay, Boqueron | | This paper |
| Hap_41 | T.garciabesi | -22 | -60,95 | Paraguay, Boqueron | | This paper |
| Hap_42 | T.garciabesi | -29 | -63 | Argentina, Santigo Estero, Aguirre | | This paper |
| Hap_42 | T.garciabesi | -28,985 | -63,450556 | Argentina, Santigo Estero, Salavina | | This paper |
| Hap_43 | T.garciabesi | -24,1833333 | -62,88333 | Argentina, Salta, Rivadavia | | This paper |
| Hap_44 | T.sordida.s.l_1 | -28,985 | -63,450556 | Argentina, Santigo Estero, Salavina | | This paper |
| Hap_45 | T.rosai | -28 | -57,587296 | Argentina, Corrientes, San Miguel. Colony. | T. sordida | Justi et al. 2014 |
| Hap_46 | T.rosai | -23,914028 | -56,701555 | Paraguay, San Pedro, Nueva Germania | | This paper |
| Hap_46 | T.rosai | -28 | -59 | Argentina, Corrientes, San Luis del Palmar. | | This paper |
| Hap_47 | T.rosai | -25,533195 | -57,079496 | Paraguay, Paraguarí, Paraguarí, Naranjo | | This paper |
| Hap_48 | T.rosai | -28 | -64 | Argentina, Santiago del Estero (Not exact locality) | | This paper |
| Hap_48 | T.rosai | -28,229857 | -61 | Argentina, Santa Fe, El nochero | | This paper |
| Hap_48 | T.rosai | -26,96 | -60,2 | Argentina, Chaco, Guemes, Colonia Aborigen | | This paper |
| Hap_48 | T.rosai | -26 | -60 | Argentina, Chaco, Guemes, Paraje El Colchon | | This paper |
| Hap_48 | T.rosai | -26,804658 | -59,997436 | Argentina, Chaco, Veinticinco de Mayo, El Triangulo | | This paper |
| Hap_48 | T.rosai | -24,93945 | -59,029187 | Argentina, Formosa, Patiño, General Manuel Belgrano | | This paper |
| Hap_48 | T.rosai | -25,64 | -60,93 | Argentina, Chaco, Guemes, La Esperanza | | This paper |
| Hap_48 | T.rosai | -26 | -60,26 | Argentina, Chaco, Maipu, La Matanza | | This paper |
| Hap_48 | T.rosai | -32 | -68 | Argentina, Mendoza, Reserva Natural Bosques Telteca | | This paper |
| Hap_48 | T.rosai | -27,78 | -64,26 | Argentina, Santiago del Estero (Not exact locality) | | This paper |
| Hap_48 | T.rosai | -24 | -57,083333 | Paraguay, San Pedro, San Pedro, Yatebu Rugua | | This paper |
| Hap_48 | T.rosai | -26 | -56,988926 | Paraguay, Paraguarí, Escobar, Chircal | | This paper |
| Hap_48 | T.rosai | -24 | -59,789786 | Paraguay, Presidente Hayes, Teniente Esteban Martínez, Tte. Esteban Martínez | | This paper |
| Hap_48 | T.rosai | -26,4 | -60,43 | Argentina, Chaco, Maipu, Tres Isletas | | This paper |
| Hap_48 | T.rosai | -27 | -58,83 | Argentina, Corrientes (Not exact locality) | | This paper |
| Hap_49 | T.rosai | -26 | -57 | Paraguay, Paraguarí, Escobar, Chircal | | This paper |
| Hap_49 | T.rosai | -26 | -57 | Paraguay, Paraguarí, Paraguarí, Mbatovi | | This paper |
| Hap_50 | T.sordida.La.Paz | -14,717401 | -68,413711 | Bolivia, La Paz, Apolo | T. guasayana | **Unpublished Belintani** |
| Hap_51 | T.sordida.La.Paz | -17 | -67,136336 | Bolivia, La Paz, Inquisivi | | This paper |
| Hap_52 | T.sordida.La.Paz | -17,796675 | -63,080339 | Bolivia, Santa Cruz. Colony. (Not exact locality) | T. guasayana | Justi et al. 2014 |
| Hap_52 | T.sordida.La.Paz | -17 | -66 | Bolivia, Cochabamba, chaco Tita. Colony. (Not exact coordinates | T. guasayana | Justi et al. 2014 |
| Hap_53 | T.sordida.s.l_2 | -17 | -66 | Bolivia, Cochabamba, Quillacollo, Cotapachi | T. sordida | Walleckx et al. 2011 |
| Hap_54 | T.sordida.s.l_2 | -17 | -66 | Bolivia, Cochabamba (Not exact coordinates) | T. sordida | Lyman et al. 1999 |
| Hap_55 | T.sordida.s.l_2 | -23 | -60 | Paraguay, Boqueron, Tiberia | | This paper |
| Hap_56 | T.sordida.s.l_2 | -17 | -68 | Bolivia, La Paz, Murillo, Mecapaca, Aucani | T. sordida | Walleckx et al. 2011 |
