## Supplementary_table_2 for "Reconstructing Sordida subcomplex (Hemiptera, Reduviidae, Triatominae) phylogeny across species distribution range"

| Haplotipo | Acc. Number |
|---|---|
| Hap_1 | **KC249290** |
| Hap_2 | **KC249287** |
| Hap_3 | **PP972075** |
| Hap_4 | **PP972076** |
| Hap_5 | **PP972083** |
| Hap_6 | **PP972082** |
| Hap_7 | **PP972084** |
| Hap_8 | **PP972087** |
| Hap_9 | **PP972088** |
| Hap_10 | **PP972085** |
| Hap_11 | **PP972086** |
| Hap_12 | **PP972077** |
| Hap_13 | **PP972080** |
| Hap_14 | **PP972079** |
| Hap_15 | **KR822194** |
| Hap_16 | **KC608980** |
| Hap_17 | **KR822187** |
| Hap_18 | **KR822188** |
| Hap_19 | **KR822193** |
| Hap_20 | **PP972078** |
| Hap_21 | **KC249291** |
| Hap_21 | **MH054942** |
| Hap_21 | **MZ700101** |
| Hap_22 | **KR822190** |
| Hap_23 | **KR822191** |
| Hap_24 | **KC249250** |
| Hap_24 | **KR822189** |
| Hap_25 | **KR822198** |
| Hap_26 | **KR822197** |
| Hap_27 | **PP972081** |
| Hap_28 | **KR822192** |
| Hap_29 | **KR822195** |
| Hap_30 | **KR822196** |
| Hap_31 | **KR822199** |
| Hap_32 | **MH054941** |
| Hap_32 | **KC249289** |
| Hap_32 | **KC249292** |
| Hap_32 | **KC249294** |
| Hap_32 | **MH054940** |
| Hap_32 | **MZ700100** |
| Hap_32 | **KR822185** |
| Hap_33 | **PP972096** |
| Hap_34 | **PP972100** |
| Hap_35 | **KC249249** |
| Hap_35 | **MH054943** |
| Hap_36 | **PP972099** |
| Hap_37 | **PP972097** |
| Hap_38 | **PP972098** |
| Hap_39 | **KC249293** |
| Hap_40 | **PP972104** |
| Hap_41 | **PP972103** |
| Hap_42 | **PP972101** |
| Hap_43 | **PP972102** |
| Hap_44 | **PP972090** |
| Hap_45 | **KC249295** |
| Hap_46 | **PP972091** |
| Hap_47 | **PP972092** |
| Hap_48 | **PP972094** |
| Hap_49 | **PP972093** |
| Hap_50 | **MH054944** |
| Hap_51 | **PP972095** |
| Hap_52 | **KC249252** |
| Hap_52 | **KC249253** |
| Hap_53 | **HQ333243** |
| Hap_54 | **AF045730** |
| Hap_55 | **PP972089** |
| Hap_56 | **HQ333242** |
