## Supplementary_table_3 for "Reconstructing Sordida subcomplex (Hemiptera, Reduviidae, Triatominae) phylogeny across species distribution range"

| | Hap_1 | Hap_2 | Hap_3 | Hap_4 | Hap_5 | Hap_6 | Hap_7 | Hap_8 | Hap_9 | Hap_10 | Hap_11 | Hap_12 | Hap_13 | Hap_14 | Hap_15 | Hap_16 | Hap_17 | Hap_18 | Hap_19 | Hap_20 | Hap_21 | Hap_22 | Hap_23 | Hap_24 | Hap_25 | Hap_26 | Hap_27 | Hap_28 | Hap_29 | Hap_30 | Hap_31 | Hap_32 | Hap_33 | Hap_34 | Hap_35 | Hap_36 | Hap_37 | Hap_38 | Hap_39 | Hap_40 | Hap_41 | Hap_42 | Hap_43 | Hap_44 | Hap_45 | Hap_46 | Hap_47 | Hap_48 | Hap_49 | Hap_50 | Hap_51 | Hap_52 | Hap_53 | Hap_54 | Hap_55 | Hap_56 |
|---|---|---|---|---|---|---|---|---|---|---|---|---|---|---|---|---|---|---|---|---|---|---|---|---|---|---|---|---|---|---|---|---|---|---|---|---|---|---|---|---|---|---|---|---|---|---|---|---|---|---|---|---|---|---|---|---|
| Hap_1 | 0 | 0,1598809325 | 0,1765670684 | 0,1713100015 | 0,2407400547 | 0,2343297105 | 0,228000512 | 0,2464763328 | 0,228000512 | 0,2407400547 | 0,2343297105 | 0,2407400547 | 0,2538126948 | 0,2604794602 | 0,2596349869 | 0,2538126948 | 0,2596349869 | 0,2538126948 | 0,2472336525 | 0,2472336525 | 0,2407400547 | 0,2472336525 | 0,2400246304 | 0,2407400547 | 0,2596349869 | 0,2538126948 | 0,2472336525 | 0,2530123736 | 0,2530123736 | 0,2407400547 | 0,2407400547 | 0,2472336525 | 0,2350493418 | 0,2286798289 | 0,2350493418 | 0,2415010442 | 0,2343297105 | 0,2286798289 | 0,2294034427 | 0,2230742443 | 0,2294034427 | 0,2415010442 | 0,2423073847 | 0,210651245 | 0,2223904378 | 0,2286798289 | 0,2294034427 | 0,2415010442 | 0,2350493418 | 0,2217504307 | 0,2286798289 | 0,2223904378 | 0,2350493418 | 0,2286798289 | 0,2286798289 | 0,2223904378 |
| Hap_2 | 0,1598809325 | 0 | 0,06861441562 | 0,06401423721 | 0,1666587045 | 0,1440559862 | 0,1552296355 | 0,1660916835 | 0,1552296355 | 0,1666587045 | 0,1609115148 | 0,1552296355 | 0,1385614254 | 0,1496115989 | 0,1491409077 | 0,1440559862 | 0,1491409077 | 0,1440559862 | 0,1385614254 | 0,1385614254 | 0,1331265892 | 0,1385614254 | 0,1381514543 | 0,1331265892 | 0,1491409077 | 0,1440559862 | 0,1331265892 | 0,1436159896 | 0,1436159896 | 0,1331265892 | 0,1331265892 | 0,1385614254 | 0,1221072971 | 0,1381514543 | 0,1381514543 | 0,1381514543 | 0,1377778862 | 0,1327459963 | 0,1385614254 | 0,1496115989 | 0,1552296355 | 0,1436159896 | 0,1331265892 | 0,1277501934 | 0,1491409077 | 0,1436159896 | 0,1552296355 | 0,154727558 | 0,1491409077 | 0,1385614254 | 0,1390075642 | 0,1445324823 | 0,1281375709 | 0,1445324823 | 0,1445324823 | 0,1390075642 |
| Hap_3 | 0,1765670684 | 0,06861441562 | 0 | 0,004305726711 | 0,1777201146 | 0,1777201146 | 0,1777201146 | 0,1889588925 | 0,1777201146 | 0,189625439 | 0,1836373434 | 0,2018211655 | 0,1956861192 | 0,1956861192 | 0,1949830618 | 0,189625439 | 0,2010806983 | 0,2018211655 | 0,1956861192 | 0,1956861192 | 0,189625439 | 0,1956861192 | 0,1949830618 | 0,189625439 | 0,2072536163 | 0,2018211655 | 0,189625439 | 0,2010806983 | 0,2010806983 | 0,189625439 | 0,189625439 | 0,1956861192 | 0,1324014903 | 0,1377778862 | 0,1377778862 | 0,1377778862 | 0,1374409699 | 0,1324014903 | 0,1381514543 | 0,1381514543 | 0,1436159896 | 0,1224309943 | 0,1327459963 | 0,1273983517 | 0,1655628118 | 0,1598809325 | 0,1660916835 | 0,1713100015 | 0,1771240205 | 0,1221072971 | 0,1277501934 | 0,1224309943 | 0,1496115989 | 0,1552296355 | 0,1552296355 | 0,1496115989 |
| Hap_4 | 0,1713100015 | 0,06401423721 | 0,004305726711 | 0 | 0,1724727235 | 0,1724727235 | 0,1724727235 | 0,1836373434 | 0,1724727235 | 0,1843075956 | 0,1783551443 | 0,1964294014 | 0,1903317648 | 0,1903317648 | 0,189625439 | 0,1843075956 | 0,1956861192 | 0,1964294014 | 0,1903317648 | 0,1903317648 | 0,1843075956 | 0,1903317648 | 0,189625439 | 0,1843075956 | 0,2018211655 | 0,1964294014 | 0,1843075956 | 0,1956861192 | 0,1956861192 | 0,1843075956 | 0,1843075956 | 0,1903317648 | 0,1273983517 | 0,1327459963 | 0,1327459963 | 0,1327459963 | 0,1324014903 | 0,1273983517 | 0,1331265892 | 0,1331265892 | 0,1385614254 | 0,1273983517 | 0,1277501934 | 0,1224309943 | 0,1603773356 | 0,154727558 | 0,1609115148 | 0,1660916835 | 0,1718720947 | 0,1171677878 | 0,1227899263 | 0,1174988716 | 0,1445324823 | 0,1501191326 | 0,1501191326 | 0,1445324823 |
| Hap_5 | 0,2407400547 | 0,1666587045 | 0,1777201146 | 0,1724727235 | 0 | 0,01746902568 | 0,03098336187 | 0,03086063937 | 0,02643787602 | 0,02643787602 | 0,02193334105 | 0,04020021272 | 0,04020021272 | 0,03098336187 | 0,03086063937 | 0,02643787602 | 0,0400352087 | 0,03557055001 | 0,03557055001 | 0,03098336187 | 0,03098336187 | 0,03098336187 | 0,03086063937 | 0,02643787602 | 0,0400352087 | 0,03557055001 | 0,03557055001 | 0,03542688115 | 0,03542688115 | 0,03557055001 | 0,03557055001 | 0,03098336187 | 0,1191679989 | 0,1300379923 | 0,1191679989 | 0,1300379923 | 0,1240758271 | 0,1191679989 | 0,1142940309 | 0,1089748318 | 0,1251052629 | 0,1032936498 | 0,1037116253 | 0,09850324483 | 0,09811225627 | 0,09298400619 | 0,09334856 | 0,09811225627 | 0,09298400619 | 0,1037116253 | 0,09374533301 | 0,08861708293 | 0,0835408972 | 0,09374533301 | 0,09374533301 | 0,08861708293 |
| Hap_6 | 0,2343297105 | 0,1440559862 | 0,1777201146 | 0,1724727235 | 0,01746902568 | 0 | 0,02193334105 | 0,02185138662 | 0,01746902568 | 0,01746902568 | 0,01304421804 | 0,03098336187 | 0,02193334105 | 0,03098336187 | 0,03086063937 | 0,02643787602 | 0,03086063937 | 0,02643787602 | 0,02643787602 | 0,02193334105 | 0,02193334105 | 0,02193334105 | 0,02185138662 | 0,01746902568 | 0,03086063937 | 0,02643787602 | 0,02643787602 | 0,02633572161 | 0,02633572161 | 0,02643787602 | 0,02643787602 | 0,02193334105 | 0,1032936498 | 0,1138203544 | 0,1032936498 | 0,1138203544 | 0,1081170256 | 0,1032936498 | 0,09850324483 | 0,09334856 | 0,1089748318 | 0,09811225627 | 0,08824647492 | 0,08319592692 | 0,0779074871 | 0,07298133888 | 0,07324534976 | 0,0779074871 | 0,07298133888 | 0,09850324483 | 0,0835408972 | 0,08861708293 | 0,07354056384 | 0,0835408972 | 0,0835408972 | 0,07851572927 |
| Hap_7 | 0,228000512 | 0,1552296355 | 0,1777201146 | 0,1724727235 | 0,03098336187 | 0,02193334105 | 0 | 0,02633572161 | 0,01304421804 | 0,02193334105 | 0,01746902568 | 0,03557055001 | 0,03557055001 | 0,03557055001 | 0,03542688115 | 0,03098336187 | 0,03542688115 | 0,03098336187 | 0,03098336187 | 0,02643787602 | 0,02643787602 | 0,02643787602 | 0,02633572161 | 0,02193334105 | 0,02633572161 | 0,02193334105 | 0,03098336187 | 0,03086063937 | 0,03086063937 | 0,03098336187 | 0,03098336187 | 0,02643787602 | 0,1032936498 | 0,1138203544 | 0,1032936498 | 0,1138203544 | 0,1081170256 | 0,1032936498 | 0,09850324483 | 0,09334856 | 0,1089748318 | 0,09811225627 | 0,08824647492 | 0,08319592692 | 0,09298400619 | 0,08790782046 | 0,07819588526 | 0,09298400619 | 0,08790782046 | 0,1142940309 | 0,09892672653 | 0,1041623765 | 0,07851572927 | 0,08861708293 | 0,08861708293 | 0,0835408972 |
| Hap_8 | 0,2464763328 | 0,1660916835 | 0,1889588925 | 0,1836373434 | 0,03086063937 | 0,02185138662 | 0,02633572161 | 0 | 0,01300159807 | 0,01300159807 | 0,008634758082 | 0,04468640503 | 0,04468640503 | 0,04468640503 | 0,04452932522 | 0,0400352087 | 0,04452932522 | 0,0400352087 | 0,0400352087 | 0,03542688115 | 0,03542688115 | 0,03542688115 | 0,03531247438 | 0,03086063937 | 0,03531247438 | 0,03086063937 | 0,0400352087 | 0,03989966252 | 0,03989966252 | 0,0400352087 | 0,0400352087 | 0,03542688115 | 0,1081170256 | 0,1186994312 | 0,1081170256 | 0,1186994312 | 0,1129738612 | 0,1081170256 | 0,1032936498 | 0,1032936498 | 0,1191679989 | 0,1029086452 | 0,1032936498 | 0,09298400619 | 0,09265187525 | 0,08760132726 | 0,08790782046 | 0,09265187525 | 0,08760132726 | 0,1138203544 | 0,09850324483 | 0,1037116253 | 0,07819588526 | 0,08824647492 | 0,08824647492 | 0,08319592692 |
| Hap_9 | 0,228000512 | 0,1552296355 | 0,1777201146 | 0,1724727235 | 0,02643787602 | 0,01746902568 | 0,01304421804 | 0,01300159807 | 0 | 0,008658225006 | 0,004310371522 | 0,03098336187 | 0,03098336187 | 0,03098336187 | 0,03086063937 | 0,02643787602 | 0,03086063937 | 0,02643787602 | 0,02643787602 | 0,02193334105 | 0,02193334105 | 0,02193334105 | 0,02185138662 | 0,01746902568 | 0,02185138662 | 0,01746902568 | 0,02643787602 | 0,02633572161 | 0,02633572161 | 0,02643787602 | 0,02643787602 | 0,02193334105 | 0,09811225627 | 0,1085292997 | 0,09811225627 | 0,1085292997 | 0,1029086452 | 0,09811225627 | 0,09334856 | 0,08824647492 | 0,1037116253 | 0,09298400619 | 0,09334856 | 0,07819588526 | 0,0779074871 | 0,07298133888 | 0,07324534976 | 0,0779074871 | 0,07298133888 | 0,09850324483 | 0,0835408972 | 0,08861708293 | 0,06373632815 | 0,07354056384 | 0,07354056384 | 0,06861441562 |
| Hap_10 | 0,2407400547 | 0,1666587045 | 0,189625439 | 0,1843075956 | 0,02643787602 | 0,01746902568 | 0,02193334105 | 0,01300159807 | 0,008658225006 | 0 | 0,004310371522 | 0,04020021272 | 0,04020021272 | 0,04020021272 | 0,0400352087 | 0,03557055001 | 0,0400352087 | 0,03557055001 | 0,03557055001 | 0,03098336187 | 0,03098336187 | 0,03098336187 | 0,03086063937 | 0,02643787602 | 0,03086063937 | 0,02643787602 | 0,03557055001 | 0,03542688115 | 0,03542688115 | 0,03557055001 | 0,03557055001 | 0,03098336187 | 0,1085292997 | 0,1191679989 | 0,1085292997 | 0,1191679989 | 0,1133802321 | 0,1085292997 | 0,1037116253 | 0,09850324483 | 0,1142940309 | 0,1032936498 | 0,1037116253 | 0,08824647492 | 0,08790782046 | 0,08288265253 | 0,08319592692 | 0,08790782046 | 0,08288265253 | 0,1089748318 | 0,09374533301 | 0,09892672653 | 0,07354056384 | 0,0835408972 | 0,0835408972 | 0,07851572927 |
| Hap_11 | 0,2343297105 | 0,1609115148 | 0,1836373434 | 0,1783551443 | 0,02193334105 | 0,01304421804 | 0,01746902568 | 0,008634758082 | 0,004310371522 | 0,004310371522 | 0 | 0,03557055001 | 0,03557055001 | 0,03557055001 | 0,03542688115 | 0,03098336187 | 0,03542688115 | 0,03098336187 | 0,03098336187 | 0,02643787602 | 0,02643787602 | 0,02643787602 | 0,02633572161 | 0,02193334105 | 0,02633572161 | 0,02193334105 | 0,03098336187 | 0,03086063937 | 0,03086063937 | 0,03098336187 | 0,03098336187 | 0,02643787602 | 0,1032936498 | 0,1138203544 | 0,1032936498 | 0,1138203544 | 0,1081170256 | 0,1032936498 | 0,09850324483 | 0,09334856 | 0,1089748318 | 0,09811225627 | 0,09850324483 | 0,08319592692 | 0,08288265253 | 0,0779074871 | 0,07819588526 | 0,08288265253 | 0,0779074871 | 0,1037116253 | 0,08861708293 | 0,09374533301 | 0,06861441562 | 0,07851572927 | 0,07851572927 | 0,07354056384 |
| Hap_12 | 0,2407400547 | 0,1552296355 | 0,2018211655 | 0,1964294014 | 0,04020021272 | 0,03098336187 | 0,03557055001 | 0,04468640503 | 0,03098336187 | 0,04020021272 | 0,03557055001 | 0 | 0,03557055001 | 0,03557055001 | 0,03542688115 | 0,03098336187 | 0,03542688115 | 0,03098336187 | 0,02193334105 | 0,02643787602 | 0,01746902568 | 0,02643787602 | 0,02633572161 | 0,02193334105 | 0,03542688115 | 0,03098336187 | 0,03098336187 | 0,03086063937 | 0,03086063937 | 0,03098336187 | 0,03098336187 | 0,02643787602 | 0,1032936498 | 0,124573457 | 0,1138203544 | 0,124573457 | 0,1186994312 | 0,1138203544 | 0,1089748318 | 0,1037116253 | 0,1196704267 | 0,1085292997 | 0,09850324483 | 0,1037116253 | 0,1032936498 | 0,09811225627 | 0,09850324483 | 0,1032936498 | 0,09811225627 | 0,1142940309 | 0,1094534311 | 0,1148010757 | 0,08861708293 | 0,07851572927 | 0,07851572927 | 0,07354056384 |
| Hap_13 | 0,2538126948 | 0,1385614254 | 0,1956861192 | 0,1903317648 | 0,04020021272 | 0,02193334105 | 0,03557055001 | 0,04468640503 | 0,03098336187 | 0,04020021272 | 0,03557055001 | 0,03557055001 | 0 | 0,01746902568 | 0,01740691291 | 0,01304421804 | 0,01740691291 | 0,01304421804 | 0,01304421804 | 0,01746902568 | 0,01746902568 | 0,01746902568 | 0,01740691291 | 0,01304421804 | 0,01740691291 | 0,01304421804 | 0,01304421804 | 0,01300159807 | 0,01300159807 | 0,01304421804 | 0,01304421804 | 0,008658225006 | 0,08790782046 | 0,1085292997 | 0,09811225627 | 0,1085292997 | 0,1029086452 | 0,09811225627 | 0,08319592692 | 0,08824647492 | 0,1037116253 | 0,09298400619 | 0,08319592692 | 0,08824647492 | 0,07298133888 | 0,06810325141 | 0,06834334972 | 0,07298133888 | 0,06810325141 | 0,09334856 | 0,07851572927 | 0,0835408972 | 0,06861441562 | 0,08861708293 | 0,08861708293 | 0,0835408972 |
| Hap_14 | 0,2604794602 | 0,1496115989 | 0,1956861192 | 0,1903317648 | 0,03098336187 | 0,03098336187 | 0,03557055001 | 0,04468640503 | 0,03098336187 | 0,04020021272 | 0,03557055001 | 0,03557055001 | 0,01746902568 | 0 | 0,008634758082 | 0,004310371522 | 0,01740691291 | 0,01304421804 | 0,01304421804 | 0,01746902568 | 0,01746902568 | 0,01746902568 | 0,01740691291 | 0,01304421804 | 0,01740691291 | 0,01304421804 | 0,01304421804 | 0,01300159807 | 0,01300159807 | 0,01304421804 | 0,01304421804 | 0,008658225006 | 0,1032936498 | 0,124573457 | 0,1138203544 | 0,124573457 | 0,1186994312 | 0,1138203544 | 0,09850324483 | 0,1037116253 | 0,1196704267 | 0,09811225627 | 0,09850324483 | 0,1037116253 | 0,09298400619 | 0,08790782046 | 0,08824647492 | 0,09298400619 | 0,08790782046 | 0,1089748318 | 0,08861708293 | 0,09374533301 | 0,07851572927 | 0,09892672653 | 0,09892672653 | 0,09374533301 |
| Hap_15 | 0,2596349869 | 0,1491409077 | 0,1949830618 | 0,189625439 | 0,03086063937 | 0,03086063937 | 0,03542688115 | 0,04452932522 | 0,03086063937 | 0,0400352087 | 0,03542688115 | 0,03542688115 | 0,01740691291 | 0,008634758082 | 0 | 0,004305726711 | 0,01737333055 | 0,01300159807 | 0,01300159807 | 0,01740691291 | 0,01740691291 | 0,01740691291 | 0,01737333055 | 0,01300159807 | 0,008658225006 | 0,01300159807 | 0,01300159807 | 0,01298733751 | 0,01298733751 | 0,01300159807 | 0,01300159807 | 0,008634758082 | 0,1029086452 | 0,1240758271 | 0,1133802321 | 0,1240758271 | 0,1182649159 | 0,1133802321 | 0,09811225627 | 0,1032936498 | 0,1191679989 | 0,09775396034 | 0,09811225627 | 0,1032936498 | 0,09265187525 | 0,08760132726 | 0,08790782046 | 0,09265187525 | 0,08760132726 | 0,1085292997 | 0,08824647492 | 0,09334856 | 0,07819588526 | 0,09850324483 | 0,09850324483 | 0,09334856 |
| Hap_16 | 0,2538126948 | 0,1440559862 | 0,189625439 | 0,1843075956 | 0,02643787602 | 0,02643787602 | 0,03098336187 | 0,0400352087 | 0,02643787602 | 0,03557055001 | 0,03098336187 | 0,03098336187 | 0,01304421804 | 0,004310371522 | 0,004305726711 | 0 | 0,01300159807 | 0,008658225006 | 0,008658225006 | 0,01304421804 | 0,01304421804 | 0,01304421804 | 0,01300159807 | 0,008658225006 | 0,01300159807 | 0,008658225006 | 0,008658225006 | 0,008634758082 | 0,008634758082 | 0,008658225006 | 0,008658225006 | 0,004310371522 | 0,09811225627 | 0,1191679989 | 0,1085292997 | 0,1191679989 | 0,1133802321 | 0,1085292997 | 0,09334856 | 0,09850324483 | 0,1142940309 | 0,09298400619 | 0,09334856 | 0,09850324483 | 0,08790782046 | 0,08288265253 | 0,08319592692 | 0,08790782046 | 0,08288265253 | 0,1037116253 | 0,0835408972 | 0,08861708293 | 0,07354056384 | 0,09374533301 | 0,09374533301 | 0,08861708293 |
| Hap_17 | 0,2596349869 | 0,1491409077 | 0,2010806983 | 0,1956861192 | 0,0400352087 | 0,03086063937 | 0,03542688115 | 0,04452932522 | 0,03086063937 | 0,0400352087 | 0,03542688115 | 0,03542688115 | 0,01740691291 | 0,01740691291 | 0,01737333055 | 0,01300159807 | 0 | 0,004305726711 | 0,01300159807 | 0,01740691291 | 0,01740691291 | 0,01740691291 | 0,01737333055 | 0,01300159807 | 0,01737333055 | 0,01300159807 | 0,01300159807 | 0,01298733751 | 0,01298733751 | 0,01300159807 | 0,01300159807 | 0,008634758082 | 0,1029086452 | 0,1240758271 | 0,1133802321 | 0,1133802321 | 0,1182649159 | 0,1133802321 | 0,09811225627 | 0,1032936498 | 0,1191679989 | 0,1081170256 | 0,09811225627 | 0,1032936498 | 0,09265187525 | 0,08760132726 | 0,08288265253 | 0,08760132726 | 0,08760132726 | 0,1138203544 | 0,09850324483 | 0,1037116253 | 0,07819588526 | 0,09850324483 | 0,09850324483 | 0,09334856 |
| Hap_18 | 0,2538126948 | 0,1440559862 | 0,2018211655 | 0,1964294014 | 0,03557055001 | 0,02643787602 | 0,03098336187 | 0,0400352087 | 0,02643787602 | 0,03557055001 | 0,03098336187 | 0,03098336187 | 0,01304421804 | 0,01304421804 | 0,01300159807 | 0,008658225006 | 0,004305726711 | 0 | 0,008658225006 | 0,01304421804 | 0,01304421804 | 0,01304421804 | 0,01300159807 | 0,008658225006 | 0,01300159807 | 0,008658225006 | 0,008658225006 | 0,008634758082 | 0,008634758082 | 0,008658225006 | 0,008658225006 | 0,004310371522 | 0,09811225627 | 0,1191679989 | 0,1085292997 | 0,1085292997 | 0,1133802321 | 0,1085292997 | 0,09334856 | 0,09850324483 | 0,1142940309 | 0,1032936498 | 0,09334856 | 0,09850324483 | 0,08790782046 | 0,08288265253 | 0,08319592692 | 0,08790782046 | 0,08288265253 | 0,1089748318 | 0,09374533301 | 0,09892672653 | 0,07354056384 | 0,09374533301 | 0,09374533301 | 0,08861708293 |
| Hap_19 | 0,2472336525 | 0,1385614254 | 0,1956861192 | 0,1903317648 | 0,03557055001 | 0,02643787602 | 0,03098336187 | 0,0400352087 | 0,02643787602 | 0,03557055001 | 0,03098336187 | 0,02193334105 | 0,01304421804 | 0,01304421804 | 0,01300159807 | 0,008658225006 | 0,01300159807 | 0,008658225006 | 0 | 0,01304421804 | 0,004310371522 | 0,01304421804 | 0,01300159807 | 0,008658225006 | 0,01300159807 | 0,008658225006 | 0,008658225006 | 0,008634758082 | 0,008634758082 | 0,008658225006 | 0,008658225006 | 0,004310371522 | 0,08790782046 | 0,1085292997 | 0,09811225627 | 0,1085292997 | 0,1029086452 | 0,09811225627 | 0,08319592692 | 0,08824647492 | 0,1037116253 | 0,09298400619 | 0,08319592692 | 0,09850324483 | 0,08790782046 | 0,08288265253 | 0,08319592692 | 0,08790782046 | 0,08288265253 | 0,09850324483 | 0,09374533301 | 0,09892672653 | 0,07354056384 | 0,0835408972 | 0,0835408972 | 0,07851572927 |
| Hap_20 | 0,2472336525 | 0,1385614254 | 0,1956861192 | 0,1903317648 | 0,03098336187 | 0,02193334105 | 0,02643787602 | 0,03542688115 | 0,02193334105 | 0,03098336187 | 0,02643787602 | 0,02643787602 | 0,01746902568 | 0,01746902568 | 0,01740691291 | 0,01304421804 | 0,01740691291 | 0,01304421804 | 0,01304421804 | 0 | 0,008658225006 | 0,008658225006 | 0,008634758082 | 0,004310371522 | 0,01740691291 | 0,01304421804 | 0,01304421804 | 0,01300159807 | 0,01300159807 | 0,01304421804 | 0,01304421804 | 0,008658225006 | 0,1032936498 | 0,124573457 | 0,1138203544 | 0,124573457 | 0,1186994312 | 0,1138203544 | 0,09850324483 | 0,1037116253 | 0,1196704267 | 0,1085292997 | 0,09850324483 | 0,09334856 | 0,08288265253 | 0,0779074871 | 0,07819588526 | 0,08288265253 | 0,0779074871 | 0,1142940309 | 0,09892672653 | 0,1041623765 | 0,06861441562 | 0,08861708293 | 0,08861708293 | 0,0835408972 |
| Hap_21 | 0,2407400547 | 0,1331265892 | 0,189625439 | 0,1843075956 | 0,03098336187 | 0,02193334105 | 0,02643787602 | 0,03542688115 | 0,02193334105 | 0,03098336187 | 0,02643787602 | 0,01746902568 | 0,01746902568 | 0,01746902568 | 0,01740691291 | 0,01304421804 | 0,01740691291 | 0,01304421804 | 0,004310371522 | 0,008658225006 | 0 | 0,008658225006 | 0,008634758082 | 0,004310371522 | 0,01740691291 | 0,01304421804 | 0,01304421804 | 0,01300159807 | 0,01300159807 | 0,01304421804 | 0,01304421804 | 0,008658225006 | 0,09298400619 | 0,1138203544 | 0,1032936498 | 0,1138203544 | 0,1081170256 | 0,1032936498 | 0,08824647492 | 0,09334856 | 0,1089748318 | 0,09811225627 | 0,08824647492 | 0,09334856 | 0,08288265253 | 0,0779074871 | 0,07819588526 | 0,08288265253 | 0,0779074871 | 0,1037116253 | 0,09892672653 | 0,1041623765 | 0,06861441562 | 0,07851572927 | 0,07851572927 | 0,07354056384 |
| Hap_22 | 0,2472336525 | 0,1385614254 | 0,1956861192 | 0,1903317648 | 0,03098336187 | 0,02193334105 | 0,02643787602 | 0,03542688115 | 0,02193334105 | 0,03098336187 | 0,02643787602 | 0,02643787602 | 0,01746902568 | 0,01746902568 | 0,01740691291 | 0,01304421804 | 0,01740691291 | 0,01304421804 | 0,01304421804 | 0,008658225006 | 0,008658225006 | 0 | 0,008634758082 | 0,004310371522 | 0,01740691291 | 0,01304421804 | 0,01304421804 | 0,01300159807 | 0,01300159807 | 0,01304421804 | 0,01304421804 | 0,008658225006 | 0,1032936498 | 0,124573457 | 0,1138203544 | 0,124573457 | 0,1186994312 | 0,1138203544 | 0,09850324483 | 0,1037116253 | 0,1196704267 | 0,1085292997 | 0,09850324483 | 0,09334856 | 0,08288265253 | 0,0779074871 | 0,07819588526 | 0,08288265253 | 0,0779074871 | 0,1142940309 | 0,09892672653 | 0,1041623765 | 0,06861441562 | 0,08861708293 | 0,08861708293 | 0,0835408972 |
| Hap_23 | 0,2400246304 | 0,1381514543 | 0,1949830618 | 0,189625439 | 0,03086063937 | 0,02185138662 | 0,02633572161 | 0,03531247438 | 0,02185138662 | 0,03086063937 | 0,02633572161 | 0,02633572161 | 0,01740691291 | 0,01740691291 | 0,01737333055 | 0,01300159807 | 0,01737333055 | 0,01300159807 | 0,01300159807 | 0,008634758082 | 0,008634758082 | 0,008634758082 | 0 | 0,004305726711 | 0,01737333055 | 0,01300159807 | 0,01300159807 | 0,01298733751 | 0,01298733751 | 0,01300159807 | 0,01300159807 | 0,008634758082 | 0,1029086452 | 0,1186994312 | 0,1133802321 | 0,1240758271 | 0,1182649159 | 0,1133802321 | 0,09811225627 | 0,1032936498 | 0,1191679989 | 0,1081170256 | 0,09811225627 | 0,09298400619 | 0,08260128559 | 0,0776507501 | 0,0779074871 | 0,08260128559 | 0,0776507501 | 0,1085292997 | 0,09334856 | 0,09850324483 | 0,06834334972 | 0,08824647492 | 0,08824647492 | 0,08319592692 |
| Hap_24 | 0,2407400547 | 0,1331265892 | 0,189625439 | 0,1843075956 | 0,02643787602 | 0,01746902568 | 0,02193334105 | 0,03086063937 | 0,01746902568 | 0,02643787602 | 0,02193334105 | 0,02193334105 | 0,01304421804 | 0,01304421804 | 0,01300159807 | 0,008658225006 | 0,01300159807 | 0,008658225006 | 0,008658225006 | 0,004310371522 | 0,004310371522 | 0,004310371522 | 0,004305726711 | 0 | 0,01300159807 | 0,008658225006 | 0,008658225006 | 0,008634758082 | 0,008634758082 | 0,008658225006 | 0,008658225006 | 0,004310371522 | 0,09811225627 | 0,1191679989 | 0,1085292997 | 0,1191679989 | 0,1133802321 | 0,1085292997 | 0,09334856 | 0,09850324483 | 0,1142940309 | 0,1032936498 | 0,09334856 | 0,08824647492 | 0,0779074871 | 0,07298133888 | 0,07324534976 | 0,0779074871 | 0,07298133888 | 0,1089748318 | 0,09374533301 | 0,09892672653 | 0,06373632815 | 0,0835408972 | 0,0835408972 | 0,07851572927 |
| Hap_25 | 0,2596349869 | 0,1491409077 | 0,2072536163 | 0,2018211655 | 0,0400352087 | 0,03086063937 | 0,02633572161 | 0,03531247438 | 0,02185138662 | 0,03086063937 | 0,02633572161 | 0,03542688115 | 0,01740691291 | 0,01740691291 | 0,008658225006 | 0,01300159807 | 0,01737333055 | 0,01300159807 | 0,01300159807 | 0,01740691291 | 0,01740691291 | 0,01740691291 | 0,01737333055 | 0,01300159807 | 0 | 0,004305726711 | 0,01300159807 | 0,01298733751 | 0,01298733751 | 0,01300159807 | 0,01300159807 | 0,008634758082 | 0,1029086452 | 0,1240758271 | 0,1133802321 | 0,1240758271 | 0,1182649159 | 0,1133802321 | 0,09811225627 | 0,1032936498 | 0,1191679989 | 0,1081170256 | 0,09811225627 | 0,1032936498 | 0,09265187525 | 0,08760132726 | 0,08790782046 | 0,09265187525 | 0,08760132726 | 0,1138203544 | 0,09850324483 | 0,1037116253 | 0,07819588526 | 0,09850324483 | 0,09850324483 | 0,09334856 |
| Hap_26 | 0,2538126948 | 0,1440559862 | 0,2018211655 | 0,1964294014 | 0,03557055001 | 0,02643787602 | 0,02193334105 | 0,03086063937 | 0,01746902568 | 0,02643787602 | 0,02193334105 | 0,03098336187 | 0,01304421804 | 0,01304421804 | 0,01300159807 | 0,008658225006 | 0,01300159807 | 0,008658225006 | 0,008658225006 | 0,01304421804 | 0,01304421804 | 0,01304421804 | 0,01300159807 | 0,008658225006 | 0,004305726711 | 0 | 0,008658225006 | 0,008634758082 | 0,008634758082 | 0,008658225006 | 0,008658225006 | 0,004310371522 | 0,09811225627 | 0,1191679989 | 0,1085292997 | 0,1191679989 | 0,1133802321 | 0,1085292997 | 0,09334856 | 0,09850324483 | 0,1142940309 | 0,1032936498 | 0,09334856 | 0,09850324483 | 0,08790782046 | 0,08288265253 | 0,08319592692 | 0,08790782046 | 0,08288265253 | 0,1089748318 | 0,09374533301 | 0,09892672653 | 0,07354056384 | 0,09374533301 | 0,09374533301 | 0,08861708293 |
| Hap_27 | 0,2472336525 | 0,1331265892 | 0,189625439 | 0,1843075956 | 0,03557055001 | 0,02643787602 | 0,03098336187 | 0,0400352087 | 0,02643787602 | 0,03557055001 | 0,03098336187 | 0,03098336187 | 0,01304421804 | 0,01304421804 | 0,01300159807 | 0,008658225006 | 0,01300159807 | 0,008658225006 | 0,008658225006 | 0,01304421804 | 0,01304421804 | 0,01304421804 | 0,01300159807 | 0,008658225006 | 0,01300159807 | 0,008658225006 | 0 | 0,008634758082 | 0,008634758082 | 0,008658225006 | 0,008658225006 | 0,004310371522 | 0,08790782046 | 0,1085292997 | 0,09811225627 | 0,1085292997 | 0,1029086452 | 0,09811225627 | 0,09334856 | 0,09850324483 | 0,1142940309 | 0,1032936498 | 0,09334856 | 0,09850324483 | 0,08790782046 | 0,08288265253 | 0,08319592692 | 0,08790782046 | 0,08288265253 | 0,09850324483 | 0,09374533301 | 0,09892672653 | 0,06373632815 | 0,0835408972 | 0,0835408972 | 0,07851572927 |
| Hap_28 | 0,2530123736 | 0,1436159896 | 0,2010806983 | 0,1956861192 | 0,03542688115 | 0,02633572161 | 0,03086063937 | 0,03989966252 | 0,02633572161 | 0,03542688115 | 0,03086063937 | 0,03086063937 | 0,01300159807 | 0,01300159807 | 0,01298733751 | 0,008634758082 | 0,01298733751 | 0,008634758082 | 0,008634758082 | 0,01300159807 | 0,01300159807 | 0,01300159807 | 0,01298733751 | 0,008634758082 | 0,01298733751 | 0,008634758082 | 0,008634758082 | 0 | 0,008639484025 | 0,008634758082 | 0,008634758082 | 0,004305726711 | 0,09775396034 | 0,1186994312 | 0,1081170256 | 0,1186994312 | 0,1129738612 | 0,1081170256 | 0,09298400619 | 0,09811225627 | 0,1138203544 | 0,1029086452 | 0,09298400619 | 0,09811225627 | 0,08760132726 | 0,08260128559 | 0,08288265253 | 0,08760132726 | 0,08260128559 | 0,1085292997 | 0,09334856 | 0,09850324483 | 0,07324534976 | 0,09334856 | 0,09334856 | 0,08824647492 |
| Hap_29 | 0,2530123736 | 0,1436159896 | 0,2010806983 | 0,1956861192 | 0,03542688115 | 0,02633572161 | 0,03086063937 | 0,03989966252 | 0,02633572161 | 0,03542688115 | 0,03086063937 | 0,03086063937 | 0,01300159807 | 0,01300159807 | 0,01298733751 | 0,008634758082 | 0,01298733751 | 0,008634758082 | 0,008634758082 | 0,01300159807 | 0,01300159807 | 0,01300159807 | 0,01298733751 | 0,008634758082 | 0,01298733751 | 0,008634758082 | 0,008634758082 | 0,008639484025 | 0 | 0,008634758082 | 0,008634758082 | 0,004305726711 | 0,09775396034 | 0,1186994312 | 0,1081170256 | 0,1186994312 | 0,1129738612 | 0,1081170256 | 0,09298400619 | 0,09811225627 | 0,1138203544 | 0,1029086452 | 0,09298400619 | 0,09811225627 | 0,08760132726 | 0,08260128559 | 0,08288265253 | 0,08760132726 | 0,08260128559 | 0,1085292997 | 0,09334856 | 0,09850324483 | 0,07324534976 | 0,09334856 | 0,09334856 | 0,08824647492 |
| Hap_30 | 0,2407400547 | 0,1331265892 | 0,189625439 | 0,1843075956 | 0,03557055001 | 0,02643787602 | 0,03098336187 | 0,0400352087 | 0,02643787602 | 0,03557055001 | 0,03098336187 | 0,03098336187 | 0,01304421804 | 0,01304421804 | 0,01300159807 | 0,008658225006 | 0,01300159807 | 0,008658225006 | 0,008658225006 | 0,01304421804 | 0,01304421804 | 0,01304421804 | 0,01300159807 | 0,008658225006 | 0,01300159807 | 0,008658225006 | 0,008658225006 | 0,008634758082 | 0,008634758082 | 0 | 0,008658225006 | 0,004310371522 | 0,08790782046 | 0,1085292997 | 0,09811225627 | 0,1085292997 | 0,1029086452 | 0,09811225627 | 0,09334856 | 0,09850324483 | 0,1142940309 | 0,1032936498 | 0,09334856 | 0,09850324483 | 0,08790782046 | 0,08288265253 | 0,08319592692 | 0,08790782046 | 0,08288265253 | 0,1089748318 | 0,09374533301 | 0,09892672653 | 0,07354056384 | 0,09374533301 | 0,09374533301 | 0,08861708293 |
| Hap_31 | 0,2407400547 | 0,1331265892 | 0,189625439 | 0,1843075956 | 0,03557055001 | 0,02643787602 | 0,03098336187 | 0,0400352087 | 0,02643787602 | 0,03557055001 | 0,03098336187 | 0,03098336187 | 0,01304421804 | 0,01304421804 | 0,01300159807 | 0,008658225006 | 0,01300159807 | 0,008658225006 | 0,008658225006 | 0,01304421804 | 0,01304421804 | 0,01304421804 | 0,01300159807 | 0,008658225006 | 0,01300159807 | 0,008658225006 | 0,008658225006 | 0,008634758082 | 0,008634758082 | 0,008658225006 | 0 | 0,004310371522 | 0,09811225627 | 0,1191679989 | 0,1085292997 | 0,1191679989 | 0,1133802321 | 0,1085292997 | 0,09334856 | 0,09850324483 | 0,1142940309 | 0,1032936498 | 0,09334856 | 0,08824647492 | 0,0779074871 | 0,07298133888 | 0,07324534976 | 0,0779074871 | 0,07298133888 | 0,09850324483 | 0,0835408972 | 0,08861708293 | 0,06373632815 | 0,0835408972 | 0,0835408972 | 0,07851572927 |
| Hap_32 | 0,2472336525 | 0,1385614254 | 0,1956861192 | 0,1903317648 | 0,03098336187 | 0,02193334105 | 0,02643787602 | 0,03542688115 | 0,02193334105 | 0,03098336187 | 0,02643787602 | 0,02643787602 | 0,008658225006 | 0,008658225006 | 0,008634758082 | 0,004310371522 | 0,008634758082 | 0,004310371522 | 0,004310371522 | 0,008658225006 | 0,008658225006 | 0,008658225006 | 0,008634758082 | 0,004310371522 | 0,008634758082 | 0,004310371522 | 0,004310371522 | 0,004305726711 | 0,004305726711 | 0,004310371522 | 0,004310371522 | 0 | 0,09298400619 | 0,1138203544 | 0,1032936498 | 0,1138203544 | 0,1081170256 | 0,1032936498 | 0,08824647492 | 0,09334856 | 0,1089748318 | 0,09811225627 | 0,08824647492 | 0,09334856 | 0,08288265253 | 0,0779074871 | 0,07819588526 | 0,08288265253 | 0,0779074871 | 0,1037116253 | 0,08861708293 | 0,09374533301 | 0,06861441562 | 0,08861708293 | 0,08861708293 | 0,0835408972 |
| Hap_33 | 0,2350493418 | 0,1221072971 | 0,1324014903 | 0,1273983517 | 0,1191679989 | 0,1032936498 | 0,1032936498 | 0,1081170256 | 0,09811225627 | 0,1085292997 | 0,1032936498 | 0,1032936498 | 0,08790782046 | 0,1032936498 | 0,1029086452 | 0,09811225627 | 0,1029086452 | 0,09811225627 | 0,08790782046 | 0,1032936498 | 0,09298400619 | 0,1032936498 | 0,1029086452 | 0,09811225627 | 0,1029086452 | 0,09811225627 | 0,08790782046 | 0,09775396034 | 0,09775396034 | 0,08790782046 | 0,09811225627 | 0,09298400619 | 0 | 0,01746902568 | 0,01746902568 | 0,01746902568 | 0,02185138662 | 0,01746902568 | 0,04938127521 | 0,04468640503 | 0,05890537269 | 0,04920225643 | 0,0400352087 | 0,06861441562 | 0,1081170256 | 0,1029086452 | 0,09811225627 | 0,1081170256 | 0,1029086452 | 0,0779074871 | 0,08319592692 | 0,08824647492 | 0,08824647492 | 0,08824647492 | 0,08824647492 | 0,08319592692 |
| Hap_34 | 0,2286798289 | 0,1381514543 | 0,1377778862 | 0,1327459963 | 0,1300379923 | 0,1138203544 | 0,1138203544 | 0,1186994312 | 0,1085292997 | 0,1191679989 | 0,1138203544 | 0,124573457 | 0,1085292997 | 0,124573457 | 0,1240758271 | 0,1191679989 | 0,1240758271 | 0,1191679989 | 0,1085292997 | 0,124573457 | 0,1138203544 | 0,124573457 | 0,1186994312 | 0,1191679989 | 0,1240758271 | 0,1191679989 | 0,1085292997 | 0,1186994312 | 0,1186994312 | 0,1085292997 | 0,1191679989 | 0,1138203544 | 0,01746902568 | 0 | 0,008658225006 | 0,008658225006 | 0,01300159807 | 0,008658225006 | 0,04938127521 | 0,04468640503 | 0,04938127521 | 0,03989966252 | 0,03086063937 | 0,07851572927 | 0,1186994312 | 0,1133802321 | 0,1085292997 | 0,1081170256 | 0,1029086452 | 0,06810325141 | 0,07324534976 | 0,07819588526 | 0,08824647492 | 0,08824647492 | 0,08824647492 | 0,08319592692 |
| Hap_35 | 0,2350493418 | 0,1381514543 | 0,1377778862 | 0,1327459963 | 0,1191679989 | 0,1032936498 | 0,1032936498 | 0,1081170256 | 0,09811225627 | 0,1085292997 | 0,1032936498 | 0,1138203544 | 0,09811225627 | 0,1138203544 | 0,1133802321 | 0,1085292997 | 0,1133802321 | 0,1085292997 | 0,09811225627 | 0,1138203544 | 0,1032936498 | 0,1138203544 | 0,1133802321 | 0,1085292997 | 0,1133802321 | 0,1085292997 | 0,09811225627 | 0,1081170256 | 0,1081170256 | 0,09811225627 | 0,1085292997 | 0,1032936498 | 0,01746902568 | 0,008658225006 | 0 | 0,008658225006 | 0,01300159807 | 0,008658225006 | 0,04938127521 | 0,04468640503 | 0,04938127521 | 0,03989966252 | 0,03086063937 | 0,07851572927 | 0,1133802321 | 0,1081170256 | 0,1032936498 | 0,1029086452 | 0,09775396034 | 0,0779074871 | 0,08319592692 | 0,08824647492 | 0,07819588526 | 0,07819588526 | 0,07819588526 | 0,07324534976 |
| Hap_36 | 0,2415010442 | 0,1381514543 | 0,1377778862 | 0,1327459963 | 0,1300379923 | 0,1138203544 | 0,1138203544 | 0,1186994312 | 0,1085292997 | 0,1191679989 | 0,1138203544 | 0,124573457 | 0,1085292997 | 0,124573457 | 0,1240758271 | 0,1191679989 | 0,1133802321 | 0,1085292997 | 0,1085292997 | 0,124573457 | 0,1138203544 | 0,124573457 | 0,1240758271 | 0,1191679989 | 0,1240758271 | 0,1191679989 | 0,1085292997 | 0,1186994312 | 0,1186994312 | 0,1085292997 | 0,1191679989 | 0,1138203544 | 0,01746902568 | 0,008658225006 | 0,008658225006 | 0 | 0,01300159807 | 0,008658225006 | 0,04938127521 | 0,04468640503 | 0,04938127521 | 0,03989966252 | 0,03086063937 | 0,07851572927 | 0,1186994312 | 0,1133802321 | 0,1085292997 | 0,1081170256 | 0,1029086452 | 0,0779074871 | 0,08319592692 | 0,08824647492 | 0,08824647492 | 0,08824647492 | 0,08824647492 | 0,08319592692 |
| Hap_37 | 0,2343297105 | 0,1377778862 | 0,1374409699 | 0,1324014903 | 0,1240758271 | 0,1081170256 | 0,1081170256 | 0,1129738612 | 0,1029086452 | 0,1133802321 | 0,1081170256 | 0,1186994312 | 0,1029086452 | 0,1186994312 | 0,1182649159 | 0,1133802321 | 0,1182649159 | 0,1133802321 | 0,1029086452 | 0,1186994312 | 0,1081170256 | 0,1186994312 | 0,1182649159 | 0,1133802321 | 0,1182649159 | 0,1133802321 | 0,1029086452 | 0,1129738612 | 0,1129738612 | 0,1029086452 | 0,1133802321 | 0,1081170256 | 0,02185138662 | 0,01300159807 | 0,01300159807 | 0,01300159807 | 0 | 0,004305726711 | 0,04452932522 | 0,04920225643 | 0,05391927255 | 0,04440213196 | 0,03531247438 | 0,08319592692 | 0,1129738612 | 0,1077382113 | 0,1029086452 | 0,1025568178 | 0,09742856769 | 0,07274875005 | 0,0779074871 | 0,08288265253 | 0,08288265253 | 0,08288265253 | 0,08288265253 | 0,0779074871 |
| Hap_38 | 0,2286798289 | 0,1327459963 | 0,1324014903 | 0,1273983517 | 0,1191679989 | 0,1032936498 | 0,1032936498 | 0,1081170256 | 0,09811225627 | 0,1085292997 | 0,1032936498 | 0,1138203544 | 0,09811225627 | 0,1138203544 | 0,1133802321 | 0,1085292997 | 0,1133802321 | 0,1085292997 | 0,09811225627 | 0,1138203544 | 0,1032936498 | 0,1138203544 | 0,1133802321 | 0,1085292997 | 0,1133802321 | 0,1085292997 | 0,09811225627 | 0,1081170256 | 0,1081170256 | 0,09811225627 | 0,1085292997 | 0,1032936498 | 0,01746902568 | 0,008658225006 | 0,008658225006 | 0,008658225006 | 0,004305726711 | 0 | 0,0400352087 | 0,04468640503 | 0,04938127521 | 0,03989966252 | 0,03086063937 | 0,07851572927 | 0,1081170256 | 0,1029086452 | 0,09811225627 | 0,09775396034 | 0,09265187525 | 0,06810325141 | 0,07324534976 | 0,07819588526 | 0,07819588526 | 0,07819588526 | 0,07819588526 | 0,07324534976 |
| Hap_39 | 0,2294034427 | 0,1385614254 | 0,1381514543 | 0,1331265892 | 0,1142940309 | 0,09850324483 | 0,09850324483 | 0,1032936498 | 0,09334856 | 0,1037116253 | 0,09850324483 | 0,1089748318 | 0,08319592692 | 0,09850324483 | 0,09811225627 | 0,09334856 | 0,09811225627 | 0,09334856 | 0,08319592692 | 0,09850324483 | 0,08824647492 | 0,09850324483 | 0,09811225627 | 0,09334856 | 0,09811225627 | 0,09334856 | 0,09334856 | 0,09298400619 | 0,09298400619 | 0,09334856 | 0,09334856 | 0,08824647492 | 0,04938127521 | 0,04938127521 | 0,04938127521 | 0,04938127521 | 0,04452932522 | 0,0400352087 | 0 | 0,03098336187 | 0,04487314393 | 0,03542688115 | 0,02643787602 | 0,06891623726 | 0,08790782046 | 0,08288265253 | 0,08824647492 | 0,08790782046 | 0,08288265253 | 0,06348894265 | 0,05890537269 | 0,06373632815 | 0,06373632815 | 0,07354056384 | 0,07354056384 | 0,06861441562 |
| Hap_40 | 0,2230742443 | 0,1496115989 | 0,1381514543 | 0,1331265892 | 0,1089748318 | 0,09334856 | 0,09334856 | 0,1032936498 | 0,08824647492 | 0,09850324483 | 0,09334856 | 0,1037116253 | 0,08824647492 | 0,1037116253 | 0,1032936498 | 0,09850324483 | 0,1032936498 | 0,09850324483 | 0,08824647492 | 0,1037116253 | 0,09334856 | 0,1037116253 | 0,1032936498 | 0,09850324483 | 0,1032936498 | 0,09850324483 | 0,09850324483 | 0,09811225627 | 0,09811225627 | 0,09850324483 | 0,09850324483 | 0,09334856 | 0,04468640503 | 0,04468640503 | 0,04468640503 | 0,04468640503 | 0,04920225643 | 0,04468640503 | 0,03098336187 | 0 | 0,01304421804 | 0,03086063937 | 0,02193334105 | 0,0543521008 | 0,09298400619 | 0,08790782046 | 0,09334856 | 0,09298400619 | 0,08790782046 | 0,0586812133 | 0,05412064719 | 0,05890537269 | 0,06861441562 | 0,06861441562 | 0,06861441562 | 0,06373632815 |
| Hap_41 | 0,2294034427 | 0,1552296355 | 0,1436159896 | 0,1385614254 | 0,1251052629 | 0,1089748318 | 0,1089748318 | 0,1191679989 | 0,1037116253 | 0,1142940309 | 0,1089748318 | 0,1196704267 | 0,1037116253 | 0,1196704267 | 0,1191679989 | 0,1142940309 | 0,1191679989 | 0,1142940309 | 0,1037116253 | 0,1196704267 | 0,1089748318 | 0,1196704267 | 0,1191679989 | 0,1142940309 | 0,1191679989 | 0,1142940309 | 0,1142940309 | 0,1138203544 | 0,1138203544 | 0,1142940309 | 0,1142940309 | 0,1089748318 | 0,05890537269 | 0,04938127521 | 0,04938127521 | 0,04938127521 | 0,05391927255 | 0,04938127521 | 0,04487314393 | 0,01304421804 | 0 | 0,03542688115 | 0,02643787602 | 0,05915983015 | 0,09811225627 | 0,09298400619 | 0,1089748318 | 0,1085292997 | 0,1032936498 | 0,06348894265 | 0,05890537269 | 0,06373632815 | 0,0835408972 | 0,0835408972 | 0,0835408972 | 0,07851572927 |
| Hap_42 | 0,2415010442 | 0,1436159896 | 0,1224309943 | 0,1273983517 | 0,1032936498 | 0,09811225627 | 0,09811225627 | 0,1029086452 | 0,09298400619 | 0,1032936498 | 0,09811225627 | 0,1085292997 | 0,09298400619 | 0,09811225627 | 0,09775396034 | 0,09298400619 | 0,1081170256 | 0,1032936498 | 0,09298400619 | 0,1085292997 | 0,09811225627 | 0,1085292997 | 0,1081170256 | 0,1032936498 | 0,1081170256 | 0,1032936498 | 0,1032936498 | 0,1029086452 | 0,1029086452 | 0,1032936498 | 0,1032936498 | 0,09811225627 | 0,04920225643 | 0,03989966252 | 0,03989966252 | 0,03989966252 | 0,04440213196 | 0,03989966252 | 0,03542688115 | 0,03086063937 | 0,03542688115 | 0 | 0,01740691291 | 0,05890537269 | 0,1029086452 | 0,09775396034 | 0,09811225627 | 0,1029086452 | 0,09775396034 | 0,06810325141 | 0,0586812133 | 0,06348894265 | 0,07324534976 | 0,07324534976 | 0,07324534976 | 0,06834334972 |
| Hap_43 | 0,2423073847 | 0,1331265892 | 0,1327459963 | 0,1277501934 | 0,1037116253 | 0,08824647492 | 0,08824647492 | 0,1032936498 | 0,09334856 | 0,1037116253 | 0,09850324483 | 0,09850324483 | 0,08319592692 | 0,09850324483 | 0,09811225627 | 0,09334856 | 0,09811225627 | 0,09334856 | 0,08319592692 | 0,09850324483 | 0,08824647492 | 0,09850324483 | 0,09811225627 | 0,09334856 | 0,09811225627 | 0,09334856 | 0,09334856 | 0,09298400619 | 0,09298400619 | 0,09334856 | 0,09334856 | 0,08824647492 | 0,0400352087 | 0,03086063937 | 0,03086063937 | 0,03086063937 | 0,03531247438 | 0,03086063937 | 0,02643787602 | 0,02193334105 | 0,02643787602 | 0,01740691291 | 0 | 0,05915983015 | 0,09811225627 | 0,09298400619 | 0,09850324483 | 0,09811225627 | 0,09298400619 | 0,06348894265 | 0,05890537269 | 0,06373632815 | 0,07354056384 | 0,07354056384 | 0,07354056384 | 0,06861441562 |
| Hap_44 | 0,210651245 | 0,1277501934 | 0,1273983517 | 0,1224309943 | 0,09850324483 | 0,08319592692 | 0,08319592692 | 0,09298400619 | 0,07819588526 | 0,08824647492 | 0,08319592692 | 0,1037116253 | 0,08824647492 | 0,1037116253 | 0,1032936498 | 0,09850324483 | 0,1032936498 | 0,09850324483 | 0,09850324483 | 0,09334856 | 0,09334856 | 0,09334856 | 0,09298400619 | 0,08824647492 | 0,1032936498 | 0,09850324483 | 0,09850324483 | 0,09811225627 | 0,09811225627 | 0,09850324483 | 0,08824647492 | 0,09334856 | 0,06861441562 | 0,07851572927 | 0,07851572927 | 0,07851572927 | 0,08319592692 | 0,07851572927 | 0,06891623726 | 0,0543521008 | 0,05915983015 | 0,05890537269 | 0,05915983015 | 0 | 0,04905332829 | 0,05374819847 | 0,06348894265 | 0,0779074871 | 0,07298133888 | 0,06834334972 | 0,05412064719 | 0,05890537269 | 0,05890537269 | 0,06861441562 | 0,06861441562 | 0,06373632815 |
| Hap_45 | 0,2223904378 | 0,1491409077 | 0,1655628118 | 0,1603773356 | 0,09811225627 | 0,0779074871 | 0,09298400619 | 0,09265187525 | 0,0779074871 | 0,08790782046 | 0,08288265253 | 0,1032936498 | 0,07298133888 | 0,09298400619 | 0,09265187525 | 0,08790782046 | 0,09265187525 | 0,08790782046 | 0,08790782046 | 0,08288265253 | 0,08288265253 | 0,08288265253 | 0,08260128559 | 0,0779074871 | 0,09265187525 | 0,08790782046 | 0,08790782046 | 0,08760132726 | 0,08760132726 | 0,08790782046 | 0,0779074871 | 0,08288265253 | 0,1081170256 | 0,1186994312 | 0,1133802321 | 0,1186994312 | 0,1129738612 | 0,1081170256 | 0,08790782046 | 0,09298400619 | 0,09811225627 | 0,1029086452 | 0,09811225627 | 0,04905332829 | 0 | 0,004310371522 | 0,03086063937 | 0,03098336187 | 0,02643787602 | 0,0779074871 | 0,06348894265 | 0,06834334972 | 0,06348894265 | 0,08319592692 | 0,08319592692 | 0,07819588526 |
| Hap_46 | 0,2286798289 | 0,1436159896 | 0,1598809325 | 0,154727558 | 0,09298400619 | 0,07298133888 | 0,08790782046 | 0,08760132726 | 0,07298133888 | 0,08288265253 | 0,0779074871 | 0,09811225627 | 0,06810325141 | 0,08790782046 | 0,08760132726 | 0,08288265253 | 0,08760132726 | 0,08288265253 | 0,08288265253 | 0,0779074871 | 0,0779074871 | 0,0779074871 | 0,0776507501 | 0,07298133888 | 0,08760132726 | 0,08288265253 | 0,08288265253 | 0,08260128559 | 0,08260128559 | 0,08288265253 | 0,07298133888 | 0,0779074871 | 0,1029086452 | 0,1133802321 | 0,1081170256 | 0,1133802321 | 0,1077382113 | 0,1029086452 | 0,08288265253 | 0,08790782046 | 0,09298400619 | 0,09775396034 | 0,09298400619 | 0,05374819847 | 0,004310371522 | 0 | 0,02633572161 | 0,02643787602 | 0,02193334105 | 0,07298133888 | 0,0586812133 | 0,06348894265 | 0,0586812133 | 0,07819588526 | 0,07819588526 | 0,07324534976 |
| Hap_47 | 0,2294034427 | 0,1552296355 | 0,1660916835 | 0,1609115148 | 0,09334856 | 0,07324534976 | 0,07819588526 | 0,08790782046 | 0,07324534976 | 0,08319592692 | 0,07819588526 | 0,09850324483 | 0,06834334972 | 0,08824647492 | 0,08790782046 | 0,08319592692 | 0,08288265253 | 0,08319592692 | 0,08319592692 | 0,07819588526 | 0,07819588526 | 0,07819588526 | 0,0779074871 | 0,07324534976 | 0,08790782046 | 0,08319592692 | 0,08319592692 | 0,08288265253 | 0,08288265253 | 0,08319592692 | 0,07324534976 | 0,07819588526 | 0,09811225627 | 0,1085292997 | 0,1032936498 | 0,1085292997 | 0,1029086452 | 0,09811225627 | 0,08824647492 | 0,09334856 | 0,1089748318 | 0,09811225627 | 0,09850324483 | 0,06348894265 | 0,03086063937 | 0,02633572161 | 0 | 0,01300159807 | 0,01740691291 | 0,09850324483 | 0,0835408972 | 0,08861708293 | 0,06861441562 | 0,07851572927 | 0,07851572927 | 0,07354056384 |
| Hap_48 | 0,2415010442 | 0,154727558 | 0,1713100015 | 0,1660916835 | 0,09811225627 | 0,0779074871 | 0,09298400619 | 0,09265187525 | 0,0779074871 | 0,08790782046 | 0,08288265253 | 0,1032936498 | 0,07298133888 | 0,09298400619 | 0,09265187525 | 0,08790782046 | 0,08760132726 | 0,08790782046 | 0,08790782046 | 0,08288265253 | 0,08288265253 | 0,08288265253 | 0,08260128559 | 0,0779074871 | 0,09265187525 | 0,08790782046 | 0,08790782046 | 0,08760132726 | 0,08760132726 | 0,08790782046 | 0,0779074871 | 0,08288265253 | 0,1081170256 | 0,1081170256 | 0,1029086452 | 0,1081170256 | 0,1025568178 | 0,09775396034 | 0,08790782046 | 0,09298400619 | 0,1085292997 | 0,1029086452 | 0,09811225627 | 0,0779074871 | 0,03098336187 | 0,02643787602 | 0,01300159807 | 0 | 0,004310371522 | 0,08790782046 | 0,07324534976 | 0,07819588526 | 0,06348894265 | 0,07324534976 | 0,07324534976 | 0,06834334972 |
| Hap_49 | 0,2350493418 | 0,1491409077 | 0,1771240205 | 0,1718720947 | 0,09298400619 | 0,07298133888 | 0,08790782046 | 0,08760132726 | 0,07298133888 | 0,08288265253 | 0,0779074871 | 0,09811225627 | 0,06810325141 | 0,08790782046 | 0,08760132726 | 0,08288265253 | 0,08760132726 | 0,08288265253 | 0,08288265253 | 0,0779074871 | 0,0779074871 | 0,0779074871 | 0,0776507501 | 0,07298133888 | 0,08760132726 | 0,08288265253 | 0,08288265253 | 0,08260128559 | 0,08260128559 | 0,08288265253 | 0,07298133888 | 0,0779074871 | 0,1029086452 | 0,1029086452 | 0,09775396034 | 0,1029086452 | 0,09742856769 | 0,09265187525 | 0,08288265253 | 0,08790782046 | 0,1032936498 | 0,09775396034 | 0,09298400619 | 0,07298133888 | 0,02643787602 | 0,02193334105 | 0,01740691291 | 0,004310371522 | 0 | 0,08288265253 | 0,06834334972 | 0,07324534976 | 0,0586812133 | 0,06834334972 | 0,06834334972 | 0,06348894265 |
| Hap_50 | 0,2217504307 | 0,1385614254 | 0,1221072971 | 0,1171677878 | 0,1037116253 | 0,09850324483 | 0,1142940309 | 0,1138203544 | 0,09850324483 | 0,1089748318 | 0,1037116253 | 0,1142940309 | 0,09334856 | 0,1089748318 | 0,1085292997 | 0,1037116253 | 0,1138203544 | 0,1089748318 | 0,09850324483 | 0,1142940309 | 0,1037116253 | 0,1142940309 | 0,1085292997 | 0,1089748318 | 0,1138203544 | 0,1089748318 | 0,09850324483 | 0,1085292997 | 0,1085292997 | 0,1089748318 | 0,09850324483 | 0,1037116253 | 0,0779074871 | 0,06810325141 | 0,0779074871 | 0,0779074871 | 0,07274875005 | 0,06810325141 | 0,06348894265 | 0,0586812133 | 0,06348894265 | 0,06810325141 | 0,06348894265 | 0,06834334972 | 0,0779074871 | 0,07298133888 | 0,09850324483 | 0,08790782046 | 0,08288265253 | 0 | 0,01740691291 | 0,01300159807 | 0,05890537269 | 0,05890537269 | 0,05890537269 | 0,05412064719 |
| Hap_51 | 0,2286798289 | 0,1390075642 | 0,1277501934 | 0,1227899263 | 0,09374533301 | 0,0835408972 | 0,09892672653 | 0,09850324483 | 0,0835408972 | 0,09374533301 | 0,08861708293 | 0,1094534311 | 0,07851572927 | 0,08861708293 | 0,08824647492 | 0,0835408972 | 0,09850324483 | 0,09374533301 | 0,09374533301 | 0,09892672653 | 0,09892672653 | 0,09892672653 | 0,09334856 | 0,09374533301 | 0,09850324483 | 0,09374533301 | 0,09374533301 | 0,09334856 | 0,09334856 | 0,09374533301 | 0,0835408972 | 0,08861708293 | 0,08319592692 | 0,07324534976 | 0,08319592692 | 0,08319592692 | 0,0779074871 | 0,07324534976 | 0,05890537269 | 0,05412064719 | 0,05890537269 | 0,0586812133 | 0,05890537269 | 0,05412064719 | 0,06348894265 | 0,0586812133 | 0,0835408972 | 0,07324534976 | 0,06834334972 | 0,01740691291 | 0 | 0,004310371522 | 0,0543521008 | 0,06401423721 | 0,06401423721 | 0,05915983015 |
| Hap_52 | 0,2223904378 | 0,1445324823 | 0,1224309943 | 0,1174988716 | 0,08861708293 | 0,08861708293 | 0,1041623765 | 0,1037116253 | 0,08861708293 | 0,09892672653 | 0,09374533301 | 0,1148010757 | 0,0835408972 | 0,09374533301 | 0,09334856 | 0,08861708293 | 0,1037116253 | 0,09892672653 | 0,09892672653 | 0,1041623765 | 0,1041623765 | 0,1041623765 | 0,09850324483 | 0,09892672653 | 0,1037116253 | 0,09892672653 | 0,09892672653 | 0,09850324483 | 0,09850324483 | 0,09892672653 | 0,08861708293 | 0,09374533301 | 0,08824647492 | 0,07819588526 | 0,08824647492 | 0,08824647492 | 0,08288265253 | 0,07819588526 | 0,06373632815 | 0,05890537269 | 0,06373632815 | 0,06348894265 | 0,06373632815 | 0,05890537269 | 0,06834334972 | 0,06348894265 | 0,08861708293 | 0,07819588526 | 0,07324534976 | 0,01300159807 | 0,004310371522 | 0 | 0,05915983015 | 0,06891623726 | 0,06891623726 | 0,06401423721 |
| Hap_53 | 0,2350493418 | 0,1281375709 | 0,1496115989 | 0,1445324823 | 0,0835408972 | 0,07354056384 | 0,07851572927 | 0,07819588526 | 0,06373632815 | 0,07354056384 | 0,06861441562 | 0,08861708293 | 0,06861441562 | 0,07851572927 | 0,07819588526 | 0,07354056384 | 0,07819588526 | 0,07354056384 | 0,07354056384 | 0,06861441562 | 0,06861441562 | 0,06861441562 | 0,06834334972 | 0,06373632815 | 0,07819588526 | 0,07354056384 | 0,06373632815 | 0,07324534976 | 0,07324534976 | 0,07354056384 | 0,06373632815 | 0,06861441562 | 0,08824647492 | 0,08824647492 | 0,07819588526 | 0,08824647492 | 0,08288265253 | 0,07819588526 | 0,06373632815 | 0,06861441562 | 0,0835408972 | 0,07324534976 | 0,07354056384 | 0,05890537269 | 0,06348894265 | 0,0586812133 | 0,06861441562 | 0,06348894265 | 0,0586812133 | 0,05890537269 | 0,0543521008 | 0,05915983015 | 0 | 0,02643787602 | 0,02643787602 | 0,02193334105 |
| Hap_54 | 0,2286798289 | 0,1445324823 | 0,1552296355 | 0,1501191326 | 0,09374533301 | 0,0835408972 | 0,08861708293 | 0,08824647492 | 0,07354056384 | 0,0835408972 | 0,07851572927 | 0,07851572927 | 0,08861708293 | 0,09892672653 | 0,09850324483 | 0,09374533301 | 0,09850324483 | 0,09374533301 | 0,0835408972 | 0,08861708293 | 0,07851572927 | 0,08861708293 | 0,08824647492 | 0,0835408972 | 0,09850324483 | 0,09374533301 | 0,0835408972 | 0,09334856 | 0,09334856 | 0,09374533301 | 0,0835408972 | 0,08861708293 | 0,08824647492 | 0,08824647492 | 0,07819588526 | 0,08824647492 | 0,08288265253 | 0,07819588526 | 0,07354056384 | 0,06861441562 | 0,0835408972 | 0,07324534976 | 0,07354056384 | 0,06861441562 | 0,08319592692 | 0,07819588526 | 0,07851572927 | 0,07324534976 | 0,06834334972 | 0,05890537269 | 0,06401423721 | 0,06891623726 | 0,02643787602 | 0 | 0,008658225006 | 0,004310371522 |
| Hap_55 | 0,2286798289 | 0,1445324823 | 0,1552296355 | 0,1501191326 | 0,09374533301 | 0,0835408972 | 0,08861708293 | 0,08824647492 | 0,07354056384 | 0,0835408972 | 0,07851572927 | 0,07851572927 | 0,08861708293 | 0,09892672653 | 0,09850324483 | 0,09374533301 | 0,09850324483 | 0,09374533301 | 0,0835408972 | 0,08861708293 | 0,07851572927 | 0,08861708293 | 0,08824647492 | 0,0835408972 | 0,09850324483 | 0,09374533301 | 0,0835408972 | 0,09334856 | 0,09334856 | 0,09374533301 | 0,0835408972 | 0,08861708293 | 0,08824647492 | 0,08824647492 | 0,07819588526 | 0,08824647492 | 0,08288265253 | 0,07819588526 | 0,07354056384 | 0,06861441562 | 0,0835408972 | 0,07324534976 | 0,07354056384 | 0,06861441562 | 0,08319592692 | 0,07819588526 | 0,07851572927 | 0,07324534976 | 0,06834334972 | 0,05890537269 | 0,06401423721 | 0,06891623726 | 0,02643787602 | 0,008658225006 | 0 | 0,004310371522 |
| Hap_56 | 0,2223904378 | 0,1390075642 | 0,1496115989 | 0,1445324823 | 0,08861708293 | 0,07851572927 | 0,0835408972 | 0,08319592692 | 0,06861441562 | 0,07851572927 | 0,07354056384 | 0,07354056384 | 0,0835408972 | 0,09374533301 | 0,09334856 | 0,08861708293 | 0,09334856 | 0,08861708293 | 0,07851572927 | 0,0835408972 | 0,07354056384 | 0,0835408972 | 0,08319592692 | 0,07851572927 | 0,09334856 | 0,08861708293 | 0,07851572927 | 0,08824647492 | 0,08824647492 | 0,08861708293 | 0,07851572927 | 0,0835408972 | 0,08319592692 | 0,08319592692 | 0,07324534976 | 0,08319592692 | 0,0779074871 | 0,07324534976 | 0,06861441562 | 0,06373632815 | 0,07851572927 | 0,06834334972 | 0,06861441562 | 0,06373632815 | 0,07819588526 | 0,07324534976 | 0,07354056384 | 0,06834334972 | 0,06348894265 | 0,05412064719 | 0,05915983015 | 0,06401423721 | 0,02193334105 | 0,004310371522 | 0,004310371522 | 0 |
